# Tumor cell villages define the co-dependency of tumor and microenvironment in liver cancer

**DOI:** 10.1101/2025.03.07.642107

**Authors:** Meng Liu, Maria O. Hernandez, Darko Castven, Hsin-Pei Lee, Wenqi Wu, Limin Wang, Marshonna Forgues, Jonathan M. Hernandez, Jens U. Marquardt, Lichun Ma

**Affiliations:** Cancer Data Science Laboratory, Center for Cancer Research, National Cancer Institute, Bethesda, Maryland 20892, USA; Spatial Imaging Technology Resource, Center for Cancer Research, National Cancer Institute, Bethesda, Maryland, USA; Department of Medicine I, University Medical Center, Lübeck, Germany; Laboratory of Human Carcinogenesis, Center for Cancer Research, National Cancer Institute, Bethesda, MD 20892, USA; Surgical Oncology Program, Center for Cancer Research, National Cancer Institute, Bethesda, Maryland 20892, USA; Liver Cancer Program, Center for Cancer Research, National Cancer Institute, Bethesda, Maryland 20892, USA

**Author notes:** Correspondence (L.M.) or (J.U.M.); <u>Main Contact:</u> Lichun Ma, PhD, Cancer Data Science Laboratory, Center for Cancer Research, National Cancer Institute, NIH, 41 Medlars Drive, Bldg. 41 Rm. A100B, Bethesda, MD 20892.

**Keywords:** Tumor cell village, Co-dependency, Spatial Dynamics Network, Spatial transcriptomics, Single-cell spatial transcriptomics, Liver cancer, Deep learning

## Abstract

Spatial cellular context is crucial in shaping intratumor heterogeneity. However, understanding how each tumor establishes its unique spatial landscape and what factors drive the landscape for tumor fitness remains significantly challenging. Here, we analyzed over 2 million cells from 50 tumor biospecimens using spatial single-cell imaging and single-cell RNA sequencing. We developed a deep learning-based strategy to spatially map tumor cell states and the architecture surrounding them, which we referred to as Spatial Dynamics Network (SDN). We found that different tumor cell states may be organized into distinct clusters, or ‘villages’, each supported by unique SDNs. Notably, tumor cell villages exhibited village-specific molecular co-dependencies between tumor cells and their microenvironment and were associated with patient outcomes. Perturbation of molecular co-dependencies via random spatial shuffling of the microenvironment resulted in destabilization of the corresponding villages. This study provides new insights into understanding tumor spatial landscape and its impact on tumor aggressiveness.

## INTRODUCTION

Intratumor heterogeneity (ITH) is a major hurdle for effective interventions across a broad range of cancers. It has been increasingly studied by single-cell transcriptome analysis, where each tumor may contain various transcriptomic states distinguished by unique gene expression patterns.^1–3^ In pan-cancer single-cell transcriptome studies, diverse tumor cell transcriptomic states have been defined, including cell cycle, epithelial-to-mesenchymal transition (EMT), and major histocompatibility complex (MHC) II-related states, among others.^4,5^ While these features of ITH are well observed, the mechanisms by which diverse tumor cells spatially coordinate and how each tumor establishes its unique spatial landscape remain poorly understood. Spatial transcriptomic profiling of various cancers at the spot resolution (a mixture of cells in each spot) has begun to unravel the intratumor spatial landscape, where a non-random spatial distribution of diverse transcriptome-based tumor clusters is observed within individual tumors.^6–13^ In ovarian cancer, a specific tumor cell state was found to spatially colocalize with tumor-infiltrating lymphocytes based on single-cell spatial transcriptome analysis.^14^ These observations suggests that the spatial cellular context is crucial in shaping ITH and further motivated us to understand the spatial landscape of individual tumors.

A critical challenge is how to spatially define tumor cell landscapes and identify the drivers underlying these diverse spatial architectures. Tumors from different patients evolve independently, leading to unique gene expression profiles that reflect patient-specific characteristics. Directly clustering spatially resolved tumor spots or cells within each tumor tends to capture patient-specific patterns, making it difficult to compare the landscapes across patients. Here, we introduce the concept of “villages” to define the distinct landscapes formed by tumor cells, where each village arises from the spatial coordination of diverse tumor cell transcriptomic states supported by specific local environments, termed the Spatial Dynamics Network (SDN), surrounding individual tumor cells. Tumor villages represent the smallest community within a tumor. This concept draws inspiration from human villages, which are small, close-knit communities where individuals share resources and responsibilities. Like human villages, tumor cell “villages” may use communal mechanisms to promote growth and enhance defense, reducing individual cell vulnerability. The introduction of this village concept provides insights into how tumor cells establish distinct spatial landscapes within individual patients, while also enabling comparisons of these landscapes across patients. More importantly, the defined tumor cell villages may uncover unique molecular co-dependencies among cells that are crucial for orchestrating their organization within each village, offering new opportunities for therapeutic interventions.

We used primary liver cancer (PLC) in this study due to its well-known, extensive ITH, making it an ideal model to investigate the spatial relationships of diverse tumor cell populations within individual tumors.^1,6,15–18^ We performed transcriptomic profiling of liver tumors at single-cell spatial resolution using CosMx™ SMI 1000-plex in situ spatial imaging platform.^19^ Additionally, we did single-cell RNA-sequencing (scRNA-seq) of tumors from the same patients to generate more comprehensive expression profiles of individual cells. In total, we profiled ~2.4 million cells from 50 tumor biospecimens. We defined tumor cell transcriptomic states and the SDNs surrounding individual tumor cells. We observed distinct spatial preferences of tumor cell states and the coordination of diverse tumor cells as tumor cell “villages” supported by unique SDNs. Tumor cell villages were found to be linked to patient outcomes. Notably, village-specific molecular co-dependencies between tumor cells and their microenvironment were observed, the perturbation of which led to the destabilization of the corresponding tumor cell villages. We validated our findings using 10X Visium spatial transcriptome data from 39 HCC patients and bulk transcriptome data from 674 HCC patients. This study may provide crucial insights into our understanding of diverse tumor spatial architectures and their impact on tumor aggressiveness.

## RESULTS

### Single-cell spatial characterization of tumors from primary liver cancer patients

To comprehensively capture ITH and investigate its stability within individual tumors, we designed the study to collect samples from multiple locations within each tumor from PLC patients (Figure 1A). In total, 50 samples were collected from the tumor core, tumor border, and adjacent normal tissue (Table S1). Among them, 15 samples (whole tissue section) were applied for single-cell spatial transcriptome profiling using the CosMx™ SMI 1000-plex in situ imaging platform. The remaining samples were subjected to single-cell RNA-sequencing (scRNA-seq) using the 10X Genomics droplet-based 5’ assay.^20^ In this study, we refer to the CosMx™ SMI data as ‘single-cell spatial data’ and the scRNA-seq data as ‘single-cell data’. After quality control (Methods), spatial transcriptomic profiles of 2,347,589 cells were derived (Figures 1B, S1A, and S1B). In addition, scRNA-seq data of 112,506 cells were generated (Figure 1B).^20^ When mapping individual cells to their spatial locations, we observed a strong concordance among cell type annotations, the expression of cell type-specific marker genes and protein staining, demonstrating a good representation of the tumor landscape with the single-cell spatial data (Figures 1C-1E, S1B, S1C, and Table S2).

**Figure 1.**
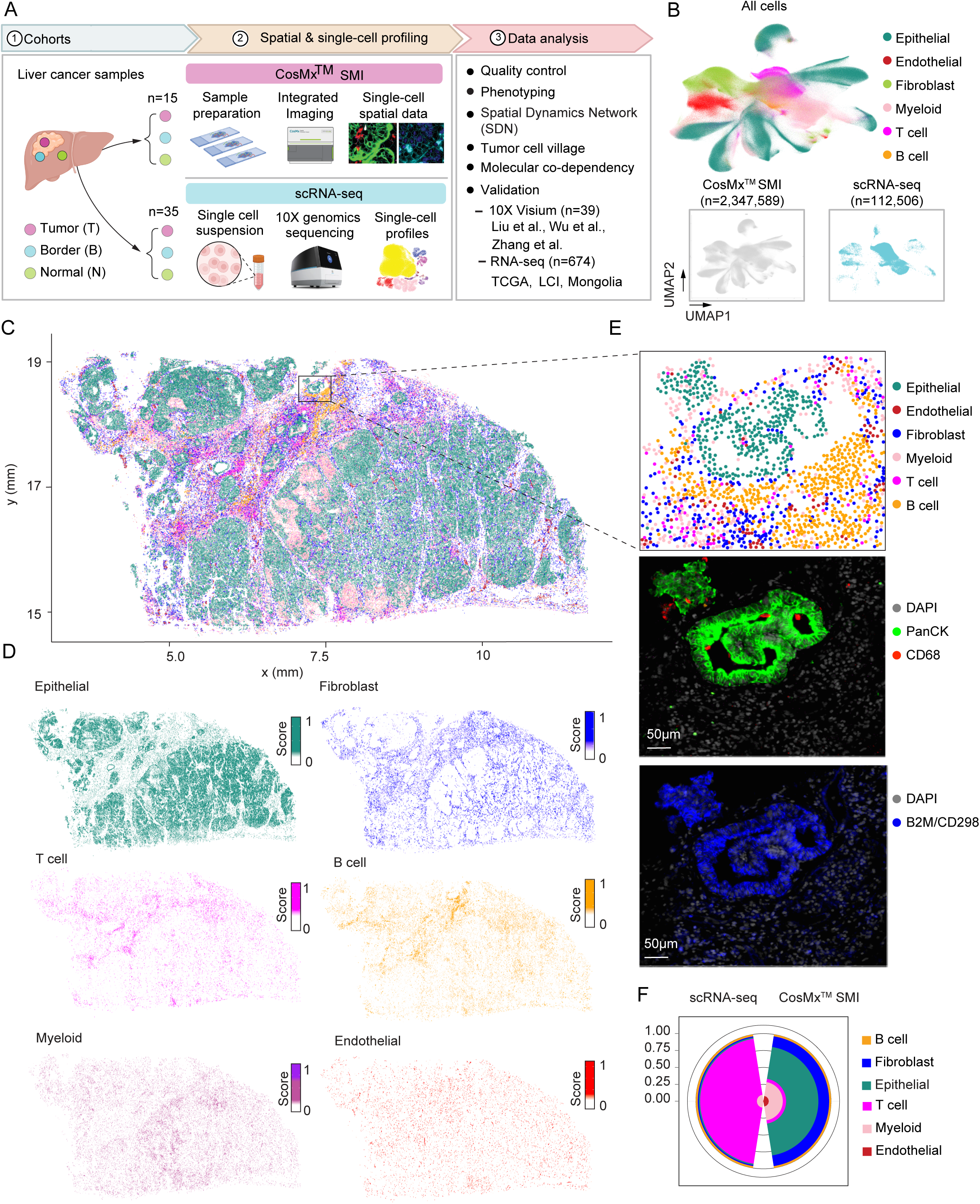
Single-cell spatial transcriptomic profiling of primary liver cancer. (A) Schematic overview of the workflow in this study. A total of 50 samples from liver cancer patients were profiled. (B) UMAP embeddings of all profiled single cells colored by cell types (n=2,460,095, top panel), and the cells colored by profiling methods (n=2,347,589, CosMx™ SMI; n=112,506, scRNA-seq, bottom panel). (C-D) A representative tumor sample (1CT) colored by cell types (C) and gene score of each cell type (D). Gene score was determined based on the average expression of marker genes specific to each cell type. (E) Cell type annotation and protein staining of a selected window in (C). CD68 (red) and Pan-cytokeratin (Pan-CK, green) represent markers for macrophages and epithelial cells. DAPI (light grey) and CD298/B2M (blue) were used for nuclei and membrane staining. Scale bars: 50µm. (F) Comparison of cell type compositions based on scRNA-seq and CosMx™ SMI from the same set of liver cancer patients in our cohort.

To further determine the quality of the single-cell spatial transcriptome data, we compared it with the scRNA-seq data from the same patients. We found strong correlations in gene expression between the two platforms, confirming a high quality of the spatial data (Figure S1D). Interestingly, we observed a striking difference in cell type composition from the two platforms. In the scRNA-seq data, T cells account for over 80% of the entire cell population (Figure 1F). By contrast, epithelial cells represent the major cell type in the single-cell spatial data, which is consistent with the histological images of the tumors.^20^ This discrepancy may be due to tissue dissociation, storage, and library preparation processes in scRNA-seq, during which immune cells may be preferentially preserved.^21^ These observations suggest that CosMx platform may offer an unbiased cellular map of liver tumors compared with scRNA-seq.

We distinguished malignant cells from non-malignant epithelial cells in the single-cell spatial data based on their transcriptomes and geographic locations in a tumor (Figure S1E). Specifically, we performed clustering analysis of all epithelial cells and mapped each cluster to its spatial location. Clusters that mapped to tumor regions were labeled as malignant cells while those mapped to adjacent normal regions were defined as epithelial cells based on histological images (see Methods). We did not infer copy number variations from the transcriptome to confirm the malignancy of epithelial cells due to a limited number of targeted genes in the single-cell spatial data. In total, 792,192 malignant cells and 1,555,397 non-malignant cells were identified from the single-cell spatial data. Malignant and non-malignant cells from the scRNA-seq data have been determined from our previous analysis.^20^

### Heterogeneous transcriptomic states in malignant cells

To understand the degree of ITH in PLC, we determined the transcriptomic states across all malignant cells from the single-cell spatial data using non-negative matrix factorization (NMF), a method has been successfully applied in defining tumor cell states based on scRNA-seq data.^4,5^ We used scRNA-seq data as a reference to increase confidence in cellular states identification (Methods). Overall, we identified 12 distinct tumor cell states across all malignant cells tested, including 9 unique states of cell cycle, stress response, immune response and locomotion, metallothionein, EMT, hepatocyte-like, cholangiocyte-like, MHC-II response, interferon response, as well as 3 mixed states of stress/metallothionein, stress/immune response and locomotion, and MHC-II/metallothionein (Figures 2A, 2B, and S2A-S2C). These states were annotated based on highly expressed genes and their functional enrichment (Figure 2B and Table S3). Immune response and locomotion (*CCL20*, *CXCL5*, *ICAM1*) represented the most prevalent cellular states in malignant cells, highlighting the interactions between tumor cells and immune cells (Figure S2D). Other frequent states included stress/metallothionein, cholangiocyte-like, and cell cycle (Figure S2D). Interestingly, we observed a small proportion of MHC II response-related tumor cells. Although MHC II molecules are primarily expressed by professional antigen-presenting cells, tumor cells can also express these molecules, and their specific expression by tumor cells has been linked to favorable outcomes in cancer patients.^22^ In contrast to malignant cells, non-malignant epithelial cells were mainly enriched in metallothionein and stress-associated programs, which are related to the functions of hepatocytes (Figure S2D). We found most of the states are similar to those identified in pan-cancer studies, suggesting commonalities of tumor cell transcriptomic states among cancer types.^4,5^ However, hepatocyte- and cholangiocyte-like states were uniquely observed in liver cancer, reflecting organ-specific cell lineages. Noticeably, each tumor comprised a mixture of different tumor cell states, indicating unique tumor cell landscape within each lesion (Figure S2E). We found that the composition of tumor cell states between the tumor border and tumor core regions of the same patient is much more similar than that among different patients, suggesting relatively stable tumor cell populations within individual patients (Figure S2E).

**Figure 2.**
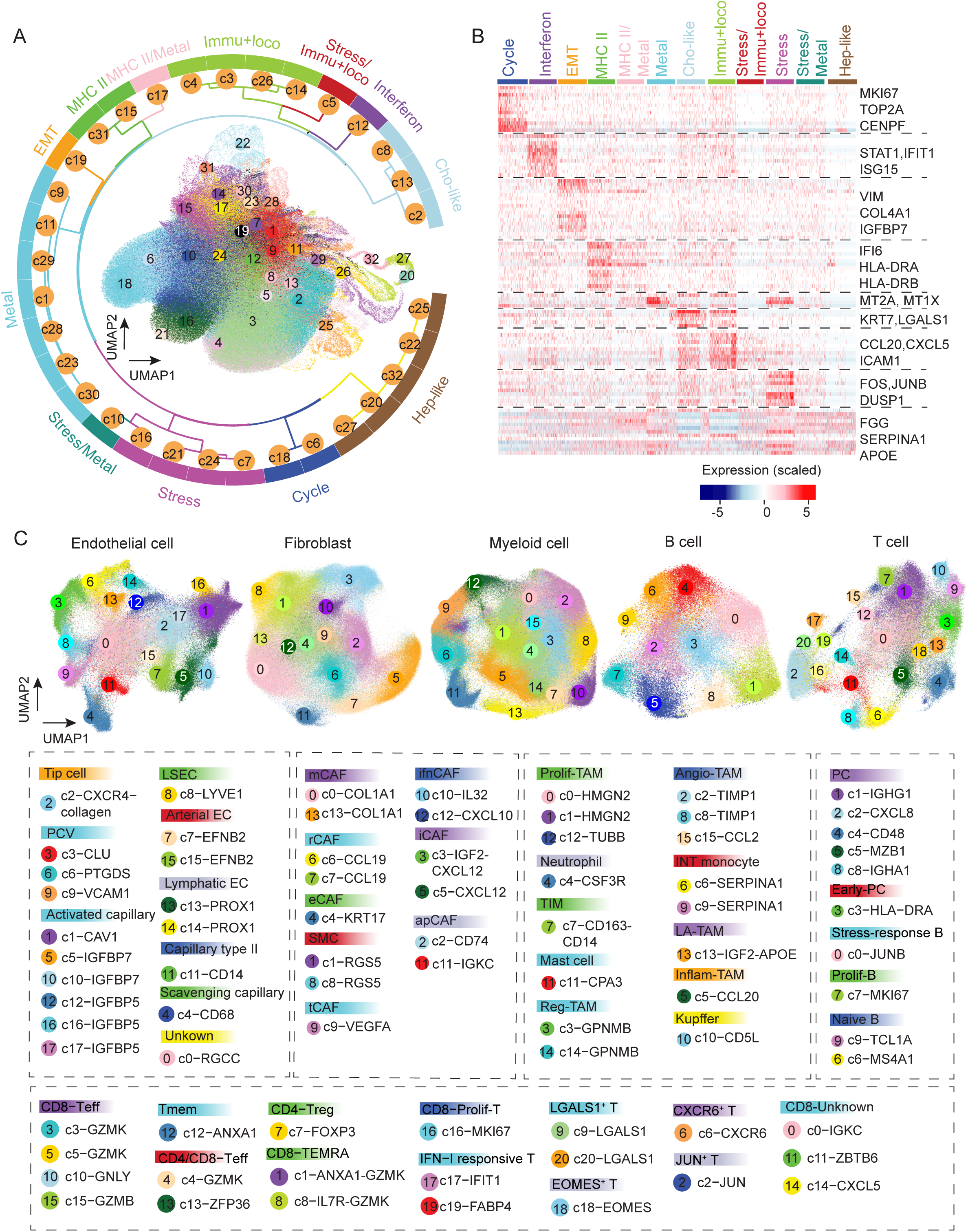
Landscape of cells in liver cancer. (A) Transcriptomic states of malignant cells. UMAP of malignant cells colored by clusters (inner plot) and a radial dendrogram indicating the hierarchical relationship of the clusters (outer dendrogram). Clusters were determined based on gene module scores of malignant cells (see Methods for details). (B) The gene expression profiles related to each tumor cell transcriptomic state in (A). Representative genes were indicated. (C) UMAP of non-malignant cells. Clusters of each non-malignant cell type were determined and annotated based on marker genes. EC, endothelial cell; PCV, post-capillary vein; LSEC, liver sinusoidal endothelial cell; CAF, cancer-associated fibroblast; mCAF, matrix CAF; rCAF, reticular-like CAF; eCAF, EMT-like CAF; SMC, smooth muscle cell; tCAF, tumour-like CAF; ifnCAF, interferon-response CAF; iCAF, inflammatory CAF; apCAF, antigen-presenting CAF; TAM, tumor-associated macrophage; Prolif-TAM, proliferating TAM; TIM, tumor infiltrating monocyte; Reg-TAM, immune-regulatory TAM; Angio-TAM, pro-angiogenic TAM; INT monocyte, intermediate monocyte; LA-TAM, lipid-associated TAM; Inflam-TAM, inflammatory cytokine-enriched TAM; PC, plasma cell; Prolif-B, proliferative B cell; Teff, effector T cell; Tmem, Memory T cell; TEMRA, recently activated effector memory or effector T cell; Treg, regulatory T cell; Prolif-T, proliferative T.

### Landscape of non-malignant cells

We performed clustering analysis of each non-malignant cell type in the single-cell spatial data to map the landscape of non-malignant cells. In total, we identified 79 clusters, comprising 18 clusters of endothelial cells, 14 clusters of fibroblasts, 16 clusters of myeloid cells, 10 clusters of B cells, and 21 clusters of T cells, reflecting a complex and heterogeneous landscape of immune and stromal cells in liver cancer (Figures 2C and S3). We further annotated these clusters into 44 cellular states based on differentially expressed genes (Figures 2C and S3). For example, in the fibroblast population, matrix fibroblasts (*COL1A1*), inflammatory fibroblasts (*IGF2*, *CXCL12*,), antigen-presenting fibroblasts (*CD74*, *IGKC*), smooth muscle cells (*RGS5*), EMT-like fibroblasts (*KRT17*), interferon-response fibroblasts (*IL32*, *CXCL10*), tumor-like fibroblasts (*VEGFA*), and reticular-like fibroblasts (*CCL19*) were determined. Similarly, different cellular states in endothelial cells, myeloid cells, and lymphoid cells were defined (Figure 2C). To determine if the non-malignant cell subtypes resolved from the single-cell spatial data and the scRNA-seq data are consistent, we compared the clusters in each major cell type derived from the two approaches. We found that most clusters from the spatial data could be well-matched to specific clusters in the scRNA-seq data, indicating that the spatial approach effectively captures major cellular information despite targeting fewer genes than scRNA-seq (Figure S4). Interestingly, we observed some unique clusters in the spatial data. For example, in myeloid cells, the *TUBB*+ proliferative tumor-associated macrophages (TAMs) were only found in the spatial data, suggesting that the single-cell spatial method may identify potential novel cell clusters (Figure S4).^23,24^ Collectively, the single-cell spatial transcriptomes resolved by CosMx offers a detailed landscape of non-malignant cells in liver cancer.

### Characterization of the surroundings of individual malignant cells by defining spatial dynamics network

A successfully evolved tumor represents a spatially well-organized ecosystem, where malignant cells actively interact with their local environment. This dynamic interplay continuously shapes tumor cell functions, fueling ITH. To uncover the roles of spatial context in driving ITH, it’s essential to delineate the surroundings of each individual malignant cell. The resolved single-cell spatial data in this study provide a unique and powerful approach for this purpose. We characterized the surroundings of each malignant cell by determining the composition of its neighbors within 40 µm, as in our previous study (Figure 3A).^25^ We further performed clustering analysis of all malignant cells based on their surroundings using the Louvain algorithm (Figure 3A). Consequently, malignant cells falling into the same cluster share similar spatial neighbors. This strategy is independent of transcriptome-based clustering, where tumor cells within a certain cluster have similar transcriptomes. We tested the stability of the malignant cell clusters by applying different radii of 20 µm, 60 µm, 80 µm, and 100 µm, where similar results were found to those derived from 40 µm, demonstrating stable clusters of malignant cells using the neighborhood information (Figures S5A and S5B). In total, 14 clusters of malignant cells (corresponding to 14 types of surrounding scenarios) with distinct surrounding environments were determined (Figures 3B, 3C, S5C, and S5D). We annotated different types of surrounding environments based on their compositions and designated each type as a Spatial Dynamics Network (SDN) around malignant cells (Figures 3A and 3C). Vascularized tumors represented a major group of SDNs (SDN-T1–SDN-T5), with endothelial cells in the surroundings of malignant cells. We observed a tumor-dominant environment (SDN-T8) in which malignant cells constitute the majority of surrounding cells. We also found four SDNs (SDN-T10–SDN-T14) with an enrichment of myeloid cells and fibroblasts, as illustrated in an example of an iCCA sample (Figures 3C and 3D). These findings highlight the diverse microenvironments surrounding malignant cells in liver cancer.

**Figure 3.**
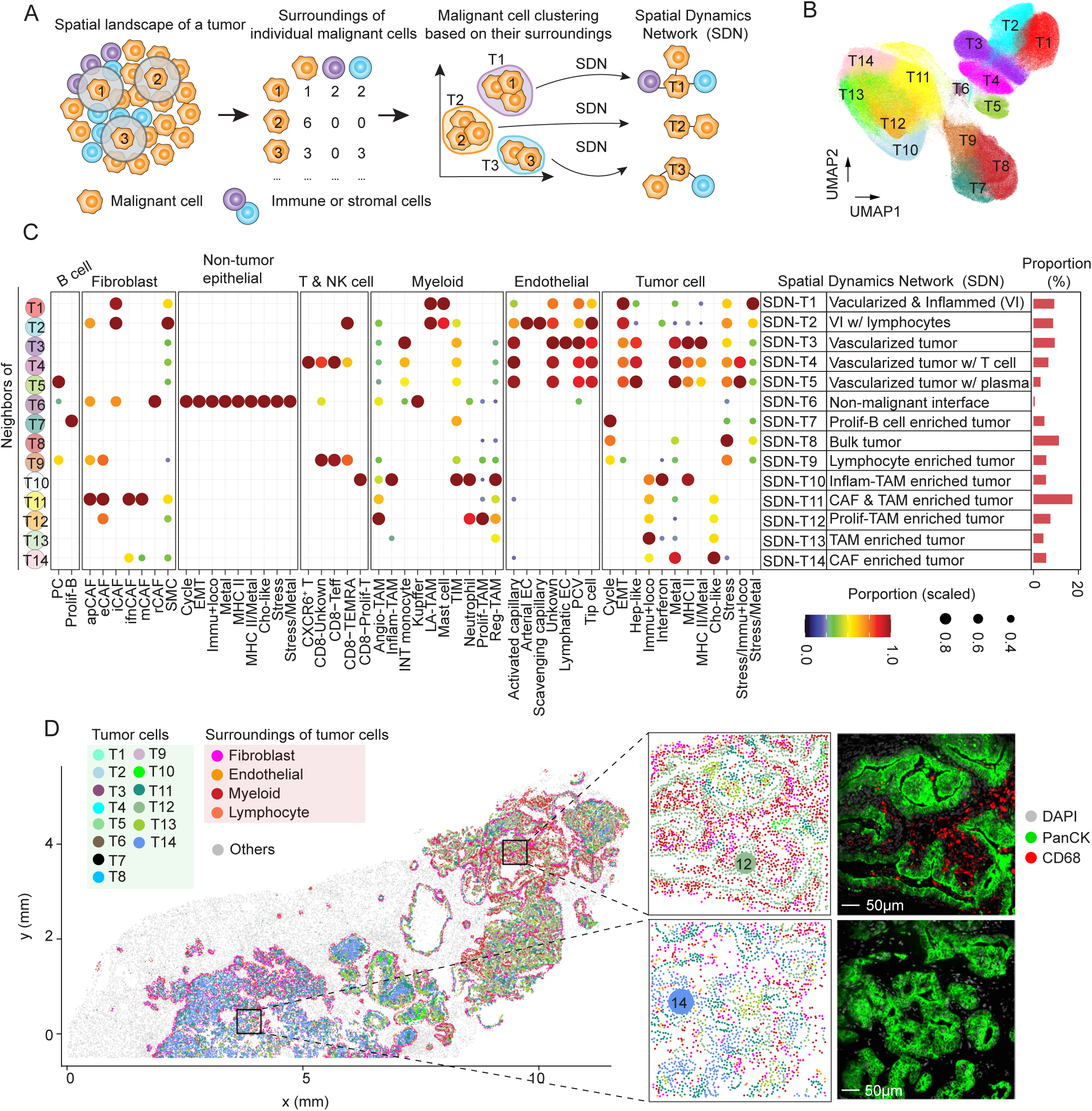
The SDNs surrounding individual malignant cells. (A) Schematic overview of SDN determination. For a targeted malignant cell, its surroundings were determined based on a 40 µm distance. Malignant cells were further characterized by the compositions of their surroundings and were subjected to clustering analysis. The average surrounding composition of each malignant cell cluster was used to define SDNs. (B) Malignant cell clusters determined based on the approach in (A). Malignant cells within the same cluster share similar surrounding features. (C) Heatmap of the surrounding compositions for each malignant cell clusters in (B). Each row of the heatmap represents the composition of the surroundings of a malignant cell cluster. Each column indicates the rescaled proportions (rescaled by column) of a cell state/subtype. (D) A representative example (1CB, left panel) of malignant cells colored by their SDNs. Malignant cells were indicated using cool colors while the surrounding non-malignant cells were shown in warm colors. Other cells are shown in grey. Two representative windows were selected with corresponding protein staining. Scale bars, 50µm.

### Tumor transcriptomic states are related to their spatial contexts

To understand the roles of spatial context in shaping ITH, we analyzed the relationship between tumor cell transcriptomic states and the SDNs. Specifically, we performed an enrichment analysis of malignant cell states on the SDNs using a random shuffling strategy (Method). Notably, tumor cell states were not randomly distributed spatially; instead, each tumor cell state was enriched in certain SDNs (Figure 4A). For example, EMT-like tumor cells were enriched in SDNs (SDN-T1–SDN-T5) that are related to vascularized tumors. Consistently, an increased density of endothelial cells along with EMT transition has been reported.^26^ In addition, we found EMT-like tumor cells were in significantly closer spatial proximity to endothelial cells compared to the tumor cell states that were enriched in completely different SDNs (Figure 4B). We observed unique populations of *IGF2*+ lipid-associated macrophages (LA-TAM) in SDN-T1 and SDN-T2, where EMT was the most strongly enriched (Figures 3C and 4A). *IGF2* has been demonstrated to be secreted by macrophages to regulate EMT.^27^ We also found inflammatory cancer-associated fibroblasts (iCAF) expressing *IGF2* and *CXCL12* in SDN-T1 and SDN-T2, suggesting a pivotal role of the local environment in regulating EMT phenotype of tumor cells (Figure 3C). In contrast to the complex local microenvironment of EMT-like tumor cells, cell cycle-related tumor cells were strongly enriched in tumor-dominant SDNs (SDN-T7–SDN-T9) with few stromal and immune cells. This was further supported by cell density analysis, with a higher density of tumor cells surrounding cell cycle-related tumor cells than others (Figure 4C). Additionally, we noticed that cell cycle- and EMT-related tumor cells were enriched in completely different SDNs. It’s well known that tumor cells going through EMT have a decreased ability to proliferate.^28^ We further demonstrate distinct local environments of the two tumor cell states. Interferons are cytokines that help trigger the immune system to eradicate pathogens or tumor cells.^29^ We observed an enrichment of interferon response-related tumor cells in SDN-T10 and SDN-T2, where myeloid cells and CD8+ T cells were found. These tumor cells expressed markers of *IFIT1* and *ISG20*, which are related to antiviral defenses.^30,31^ Noticeably, interferon response-related tumor cells were mainly found in a patient with infections of hepatitis C virus and human immunodeficiency virus in our cohort, suggesting a potential antiviral response in this patient (Figure S2E and Table S1). We also found a strong enrichment of cholangiocyte-like tumor cells and immune response and locomotion-related tumor cells in the SDNs with fibroblasts and myeloid cells (SDN-T11–SDN-T13). Ligand-receptor interaction analysis between tumor cell states and their local environments demonstrated enriched communication patterns in each state, further supporting the spatial preference of different tumor cell states (Figure S6). For example, interactions mediated by VEGF were mainly observed between EMT-related tumor cells and their surrounding environments (Figure S6).

**Figure 4.**
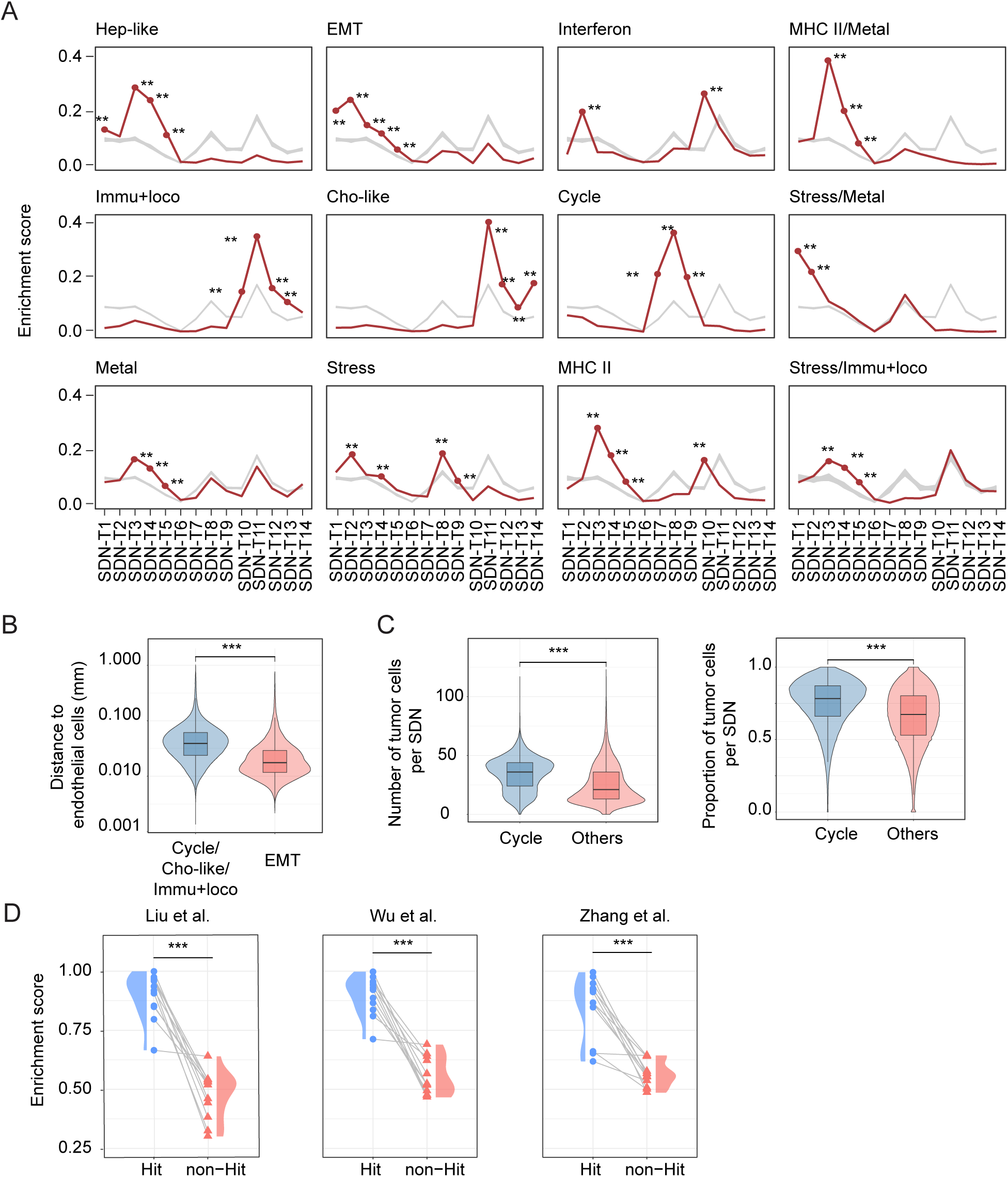
Enrichment of tumor cell states on SDNs. (A) Enrichment of each tumor cell state on SDNs. Red lines indicate observed enrichment while grey lines stand for randomized enrichment score. P-values were calculated based on the observed and randomized values (see Methods for details). **, p-value <0.01. (B) Boxplot of the distances between EMT-like malignant cells and endothelial cells as well as the distances between other malignant cells (cell cycle, cholangiocyte-like, or immune response and locomotion) and endothelial cells. Only tumor cell states with no significant enrichment in the SDN-T1–SDN-T5 were included in the comparison with EMT-like malignant cells. P-value was calculated with Student’s t-test. ****, p-value <0.001. (C) Boxplots of the number (left) and the proportion (right) of malignant cells in the surrounding (within 40 µm distance) of cell cycle-related malignant cells compared to that of other malignant cells. P-value was calculated using Student’s t-test. ***, p-value <0.001. (D) Validation of the spatial preference of tumor cell states using 10X Visium data from Liu et al. (left), Wu et al. (middle) and Zhang et al. (right). Each pair of connected dot and triangle represents a tumor cell state. For each tumor cell state, “Hit” represents the significantly associated SDNs in (A), while “non-Hit” indicates the rest of the SDNs. Violin plots of the distributions of the enrichment scores were shown. p-value was calculated with Student’s t-test. ****, p-value <0.001.

To validate the spatial preference of tumor cell states, we analyzed 39 liver tumor samples from three publicly available datasets based on the 10X Visium platform.^6,32,33^ Since 10X Visium data is at spot (55 µm in diameter) resolution, the methods designed for single-cell spatial data cannot be directly applied to determine cellular states and SDNs here. Therefore, we developed pseudo-bulk gene signatures for each tumor cell transcriptomic state and SDN by averaging the associated genes from our single-cell spatial data (Methods). We further applied these gene signatures to the three datasets and used a correlation-based approach to determine the enrichment of each tumor cell state in the SDNs (Methods). We observed highly consistent enrichment patterns with those found in the single-cell spatial data, with a much stronger enrichment of tumor cell states in SDNs that showed significant association in the single-cell spatial analysis compared to other SDNs (Figure 4D). Collectively, these observations suggest that the transcriptomic states of tumor cells are not randomly distributed within a tumor. Instead, they are closed associated with their surrounding microenvironments, underlying the potential roles of spatial context in driving ITH.

### Tumor cell villages are identified using graph attention networks

Diverse tumor cells may spatially coordinate to collectively drive tumor growth and progression, conferring survival advantages to the tumor and enhancing its resilience against interventions.^34^ We define this spatial coordination of diverse tumor cell transcriptomic states, supported by specific local environments, as tumor “villages”. Like human villages, tumor cell “villages” may use communal mechanisms to promote growth and enhance defense, reducing individual cell vulnerability.

To determine tumor cell villages, we used graph attention networks, which are deep learning neural networks specifically designed for analyzing graph-structured data.^35^ These networks leverage an attention mechanism to enhance feature learning by weighing the importance of neighboring nodes, allowing us to capture the complex spatial relationships between diverse tumor cell states. In general, malignant cells within a tumor were represented as connected graphs, where nodes indicate individual cells characterized by their cellular states and SDNs, and edges connect those nodes that are within a 40 µm distance, consistent with the SDN analysis (Figure 5A). The embeddings derived from the graph attention networks were then applied for clustering analysis to define tumor cell villages (Figure 5B). We excluded three small clusters with fewer than 5,000 cells each, accounting for less than 1% of total malignant cells. Eight distinct tumor villages were then identified (Figures 5C and S7A). Village 1 and Village 2 contain a mix of tumor cell states with a vascularized environment. By contrast, Village 4 and Village 7 are dominated by cell cycle-related tumor cells, akin to cancer germinal centers. Villages 3, 5, and 6 are enriched in cholangiocyte-like and immune response and locomotion-related tumor cells, surrounded by a microenvironment of fibroblasts and macrophages, reflecting a well-developed ecosystem of tumor cells with ample resources. Village 8 is primarily associated with immune response and locomotion. As an example, 1CB (an iCCA sample) features Villages 3, 5, and 6 in distinct geographic locations (Figure 5D). We noticed that tumor cell village compositions were much more similar across different sampling regions from the same patient compared to those from different patients, suggesting relatively stable tumor cell populations within individual patients (Figure S7B).

**Figure 5.**
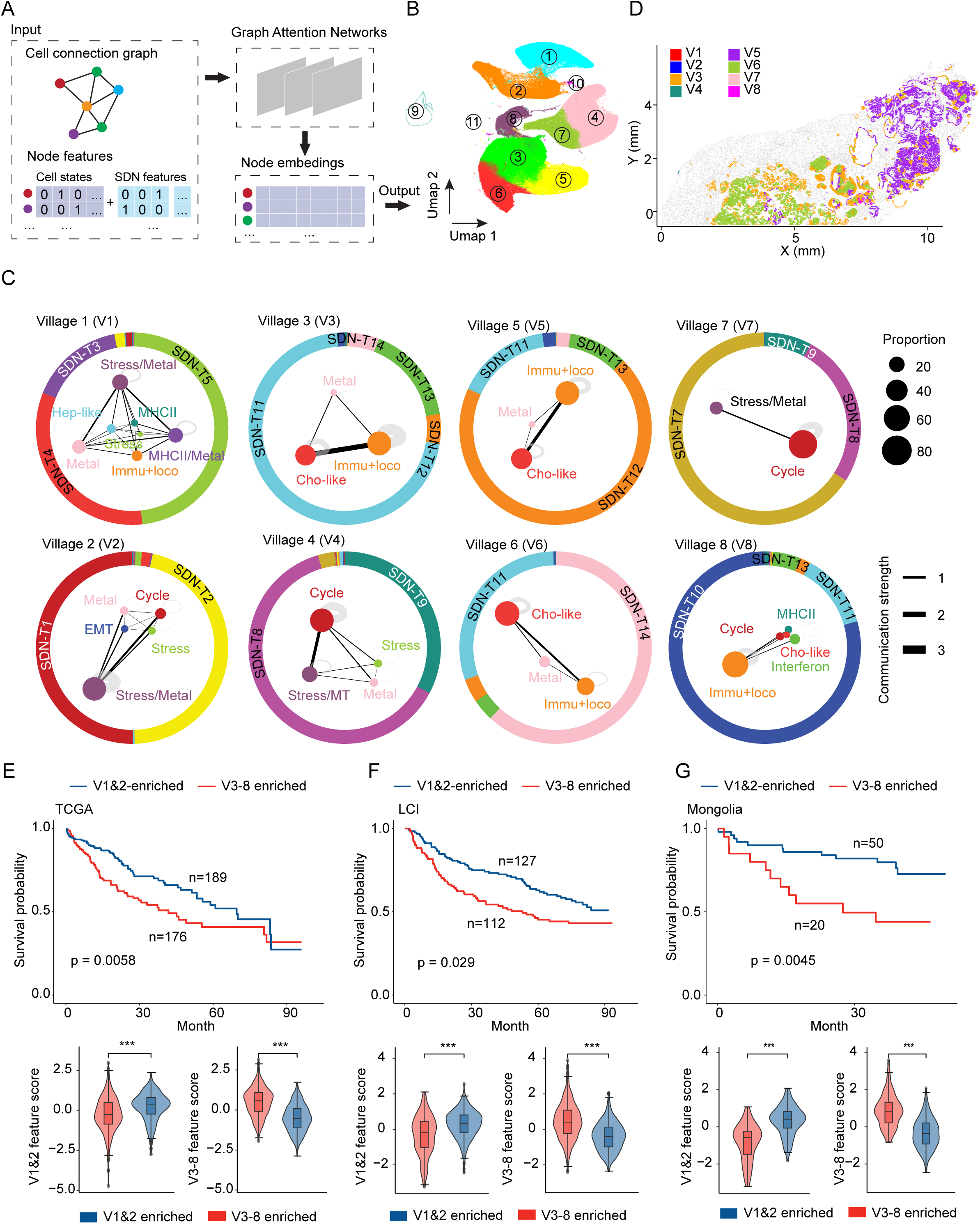
Tumor cell villages in liver cancer. (A) Schematic overview of determining tumor cell villages using graph attention network-based approach. Malignant cells were represented as connected graphs, where nodes indicate individual cells characterized by their cellular states and SDNs, and edges connect those nodes that are within a 40 µm distance. The embeddings derived from the GAT analysis were used for clustering to determine tumor cell villages. (B) UMAP of malignant cell groups based on the embeddings derived from graph attention networks in (A). (C) Village plots of different tumor cell villages. Dots represent malignant cells colored based on their cellular states. The size of each dot reflects the proportion of tumor cell states within a village. The thickness of the lines between dots indicates the strength of the communications. Outer circles indicate the SDNs associated with each village, with different colored areas represent the proportions of the SDNs. (D) A representative example (1CB) of tumor cell villages. Non-malignant cells were colored grey. (E-G) Top: Overall Survival (OS) of HCC patients enriched in V1 and V2 related features (blue) and V3-V8 related features (red) in TCGA (E, TCGA-LIHC, n=365), LCI (F, Liver Cancer Institute, n=239), and Mongolia (G, n=70) cohorts. Kaplan-Meier plots were used to show the overall survival of each patient group. p-value was calculated using the log rank test. Bottom: Village feature score of the two patient groups in the top panel. P values were calculated using Student’s t-test (***: p value<0.001).

We determined surrogate markers for tumor villages through differential gene expression analysis (Figure S7C and Table S4). The identified genes were found largely specific to each tumor village and can accurately predict the presence of their corresponding villages (Figures S7D-S7F), suggesting that these genes may effectively serve as surrogates for defining tumor villages. With these genes, we then validated the existence of the identified tumor cell villages based on spatial transcriptome data from 39 liver tumor samples generated using the 10X Visium platform (Figures S7G and S7H). Additionally, we performed survival analysis of 674 HCC patients in the TCGA, LCI (Liver Cancer Institute), and Mongolia cohorts based on hierarchical clustering of the surrogate genes (Figure S7I). We found that the patients with Villages 1 and 2-related gene signatures were consistently associated with significantly better clinical outcome compared to those with features related to Villages 3-8 (Figures 5E-5G). These analyses highlight the presence of well-coordinated tumor cell villages in liver cancer and suggest that tumor cells villages are linked to patient outcomes.

### Molecular co-dependencies linked to tumor cell villages

To uncover key factors driving cell coordination within each tumor cell village, we developed a method to evaluate the molecular co-dependencies between tumor cells and non-tumor (stromal or immune) cells. Specifically, we calculated the correlations between each pair of genes from the tumor and non-tumor cells, with an assumption that highly correlated gene pairs are strongly co-dependent (Methods). Using this method, we identified the top correlated gene pairs from each village and found that these gene pairs are village specific, with significantly higher correlation scores in their respective villages compared to other (Figure 6A; Table S5). We further evaluated the gene-pair correlations for each village using a distance-based approach, by analyzing their correlation scores at spatial distances of 40 µm, 80 µm, 160 µm, 320 µm, and extending beyond the village boundaries. Notably, a gradual decline in correlation scores was observed as the spatial separation of cells increased, underscoring the spatial specificity of the gene-pair relationships (Figure 6B). To demonstrate the importance of these gene-pair relationships to each tumor cell village, we employed a random forest model to predict village identities of the tumor cells based on the identified gene pairs. We observed a significant decline in prediction accuracy when the non-tumor cells surrounding the tumor cells within each village were randomly shuffled, compared to the accuracy achieved with the original village configurations (Figure 6C). These findings suggest that molecular co-dependencies may be crucial to maintaining the integrity of each village, and their perturbation may potentially destabilize or collapse the village structure.

**Figure 6.**
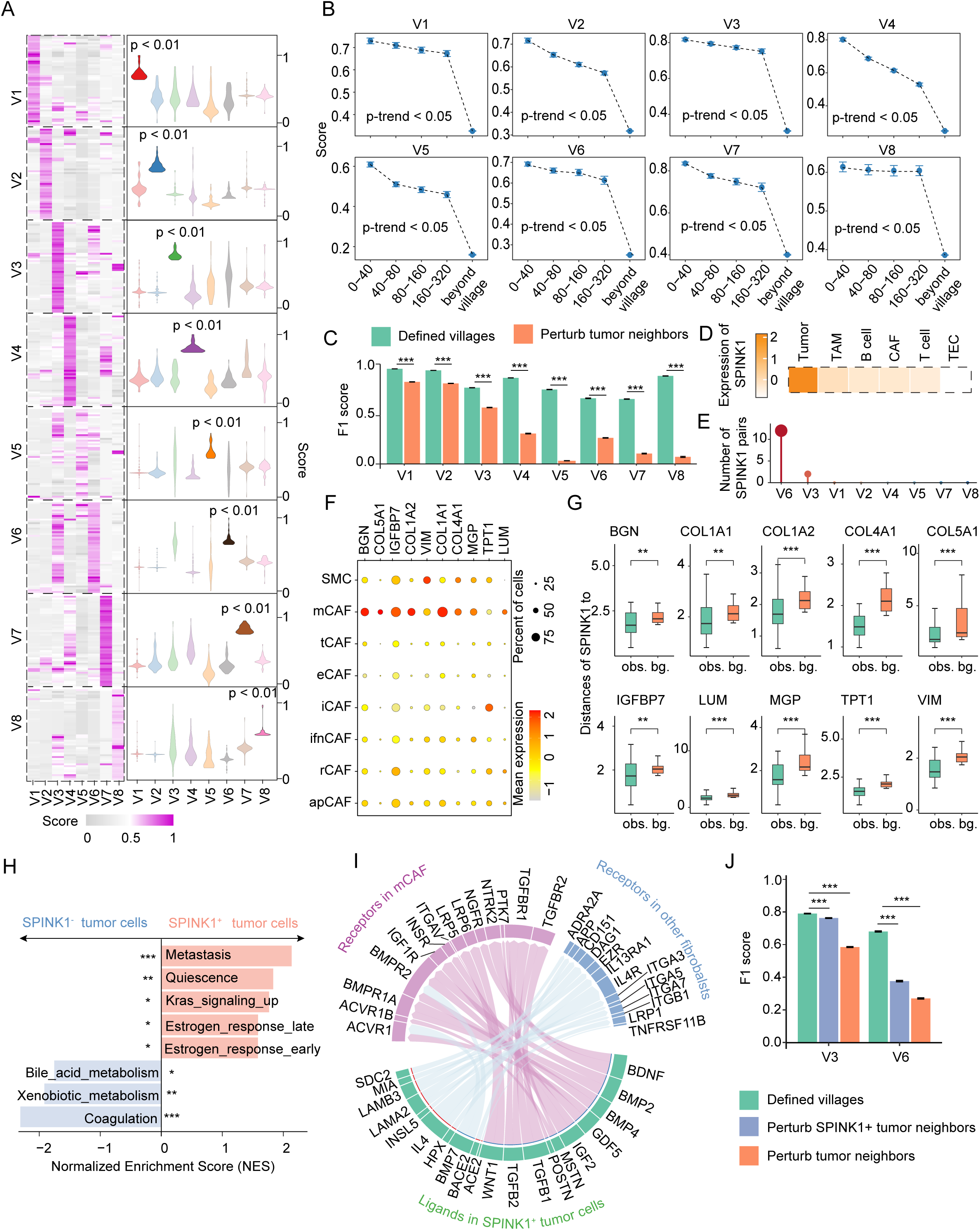
Spatial molecular co-dependencies in individual tumor cell villages. (A) Correlation scores of top gene pairs in each village. Violin plots (right) indicate the scores of the gene pairs in the heatmap for each tumor cell village. One-tailed Student’s t-test was used for statistical tests. Scores were normalized to a 0–1 scale within each village. (B) Gradient changes of correlation scores between tumor cells and their neighbors along difference spatial distances. Data are shown with mean ± standard deviation (SD). Statistical significance was assessed using the Mann-Kendall trend test. Distance, µm. Scores were normalized to a 0–1 scale within each village. (C) F1 score of the performance of the random forest model on the defined tumor villages and after perturbing the neighbors of tumor cells. Mean F1 score ± SD was shown based on the 100 iterations. (D) Expression of SPINK1 in different cell types. (E) Number of SPINK1-related gene pairs in each tumor cell village. (F) Expression of genes across different fibroblast subtypes. Color represents average gene expression, while dot size indicates the fraction of cells expressing a certain gene. (G) Distances between SPINK1^+^ and marker positive (BGN, COL1A1, COL1A2, COL4A1, COL5A1, IGFBP7, LUM, MGP, TPT1 and VIM) spots shown as observations (obs.), compared to distances between SPINK1+ and randomly selected spots as background (bg.). P-values were calculated using Student’s t-test (*: p < 0.05; **: p < 0.01; ***: p < 0.001). (H) Gene set enrichment analysis of SPINK1+ tumor cells compared with SPINK1-tumor cells. (*: p value <0.05; **: p value<0.01; ***: p value<0.001). (I) Chord plot of cell-cell communications between SPINK1+ tumor cells (green) and mCAFs (magenta) or other fibroblasts (blue). Line width represents the interaction strength. (J) F1 score of the performance of the random forest model on the defined tumor villages, perturbing the neighbors of SPINK1^+^ tumor cells, and after perturbing the neighbors of all tumor cells. Mean F1 score ± SD was shown based on the 100 iterations.

Among the top ranked molecules from tumor cells, we focused on serine protease inhibitor Kazal-type 1 (SPINK1) due to its unique expression in tumor cells (Figures 6D and S8A). SPINK1 was identified from Villages 3 and 6, with much more SPINK1-related gene pairs found in Village 6 than Village 3 (Figure 6E). Most of the SPINK1-associated genes are matrix CAFs (mCAFs)-related genes, including COL1A1, COL1A2, COL5A1, COL4A1, BGN, and MGP (Figure 6F; Table S5). This spatial relationship was further validated using Visium transcriptome data, which showed that SPINK1+ spots were significantly closer to the paired gene-enriched spots than expected by a random chance (Figures 6G). Gene set enrichment analysis suggested that SPINK1+ tumor cells were associated with metastasis-related features, while SPINK1-tumor cells were enriched in metabolism- and coagulation-related pathways (Figure 6H). Additionally, single-cell transcriptome analysis demonstrated that SPINK1+ tumor cells interacted more extensively with mCAFs than other CAF subtypes, suggesting a preferential interaction between SPINK1+ tumor cells and mCAFs (Figure 6I and Figure S8B). When the relationship of SPINK1 and its paired genes were perturbed by randomly shuffling non-tumor cells, we found that the prediction accuracy of village identities by the random forest model declined significantly compared with the original village configurations (Figure 6J). The prediction accuracy was further declined when all the identified correlated genes in these villages were perturbed (Figure 6J). These observations suggest the relationships between SPINK1+ tumor cells and mCAFs may play an important role in sustaining the integrity of Villages 3 and 6. Collectively, these findings indicate that the defined tumor cell villages may reveal unique molecular co-dependencies among cells with each village, opens new avenues for developing new treatment strategies aimed at disrupting these co-dependencies to destabilize the tumor.

## DISCUSSION

Over the past decade, transcriptomic ITH has been increasingly recognized, owing to the development of single-cell technologies. However, an essential factor in understanding ITH—the spatial context of cells—is lost in single-cell analysis. To determine the geographic landscape of diverse tumor cell transcriptomic states and their spatial relationships, we performed single-cell spatial transcriptomic profiling of liver cancer. We developed a bioinformatics method to classify the surroundings of individual malignant cells, where 14 different types of SDNs were identified, including vascularized tumor, tumor-dominant environment, and tumor-nontumor interface, among others. Interestingly, we found that each tumor cell state was preferentially associated with specific SDNs, rather than being randomly distributed throughout the tumor. The observations were further supported by spatial proximity analysis, cell density analysis, and ligand-receptor interaction analysis. These findings indicate a close link between tumor cell states and their microenvironments, suggesting a crucial role of spatial context in driving ITH. Notably, such insights would not be possible without spatial information of individual cells, highlighting the importance of single-cell spatial transcriptome approaches in advancing our understanding of ITH.

Tumor cells form diverse spatial landscapes for survival fitness. Understanding how each tumor establishes its unique spatial landscapes and what factors drive these landscapes may offer new insights into the disease development, progression as well as response to therapy. To this end, we developed a graph attention neural network-based method to identify spatially coordinated groups of tumor cells, which we termed tumor cell “villages”. This concept was inspired by human villages, which historically formed to fulfill social, economic, and environmental needs, fostering a sense of community, mutual support, security, and the ability to collectively adapt to environmental challenges. This concept differs from the cellular neighborhood methods which predominantly divide a tissue into broad regions based on major cell type compositions or general connections, where the spatial coordination of diverse tumor cell states and their specific interactions with certain SDNs are missing.^36–38^ We identified eight different types of villages among available liver tumors using graph attention neural networks. We noticed that each patient may comprise a combination of different tumor cell villages. Using surrogate markers from each tumor cell village, we found that Villages 1 and 2-related features were associated with better patient outcomes than those related to Villages 3-8. The defined tumor cell villages further allowed us to uncover unique molecular co-dependencies among cells within each village, which may play a critical role in orchestrating cell coordination and spatial organization. Interestingly, when the molecular co-dependencies were perturbed through in silico simulations, we found a significant decline in the village identity prediction accuracy by a random forest model, suggesting that the molecular co-dependencies are crucial for maintaining the integrity of a tumor cell village. Notably, SPINK1, derived from tumor cells, was spatially associated with genes expressed by mCAFs in Villages 3 and 6. SPINK1 is a serine protease inhibitor known for preventing the inappropriate activation of trypsin in the pancreas.^39^ In liver cancer, SPINK1 has been implicated in promoting tumor plasticity.^40^ However, the interactions between SPINK1+ tumor cells and other cell types, as well as their roles in the spatial organization of tumors remain poorly understood. Here, we demonstrate that SPINK1+ tumor cells have unique molecular co-dependencies with mCAFs, the perturbation of which leads to the destabilization of the corresponding tumor cell villages. These findings suggest that the defined tumor cell villages reveal village-specific molecular co-dependencies that may contribute to shaping each tumor’s unique spatial landscape, paving the way for developing new strategies for therapeutic interventions.

PLC is one of the most lethal malignancies worldwide, with a median overall survival of 6– 10 months and a 5-year survival rate of around 20%.^41,42^ Hepatocellular carcinoma (HCC) and intrahepatic cholangiocarcinoma (iCCA) are the two major clinical subtypes of PLC. Extensive ITH in both clinical subtypes has been demonstrated using single-cell approaches in our previous studies.^1,15,16^ Thus, in this study, we used PLC to model the spatial relationships among diverse malignant cells. While shared tumor cell states and tumor cell villages between HCC and iCCA were revealed in our analyses, each clinical type had specific enrichment patterns (Figures S2E and S7B). These observations align with previous studies that demonstrate both the commonality and uniqueness of the molecular features of these two subtypes.^43^

This study has limitations. First, we aimed to achieve a deep understanding of intratumor heterogeneity by performing scRNA-seq and single-cell spatial transcriptome profiling of samples from multiple locations within a tumor from seven liver cancer patients. To validate our findings, we used 3 publicly available spatial datasets including 39 liver tumor samples based on the 10x Visium platform, and 674 HCC patients with bulk transcriptome data. Second, validating the link between tumor cell states and the spatial contexts of the cells using in vitro and in vivo models may provide additional insights into tumor heterogeneity and plasticity. Future co-culture experiments or in vivo studies need to be carefully designed for this purpose. Third, we demonstrated a village concept of tumor cells in liver cancer. Extending this concept to other cancer types may help further understand diverse tumor cell spatial relationships.

## ACKNOWLEDGEMENTS

We thank Drs. Xin Wei Wang, Eytan Ruppin, and Tom Misteli for helpful comments on the manuscript; Dr. Yuuki Ohara for assistance in interpreting the histology images; the patients, families, and nurses for contribution to this study. This work was supported by grants (ZIA BC 012079 and ZIA BC 012083) from the intramural research program of the Center for Cancer Research, National Cancer Institute of the United States. JUM is supported by grants from the Wilhelm Sander Foundation (2021.089.1)

## AUTHOR CONTRIBUTIONS

L.M. developed study concept; J.U.M. directed clinical study; M.L. performed computational analysis; M.O.H. conducted experiments; D.C., H.L., W.W., L.W., M.F., J.M.H. conducted additional experiments and data analysis; M.L. and L.M. interpreted data; L.M., and M.L. wrote the manuscript with help from M.O.H. All authors read, edited, and approved the manuscript.

## METHODS

### Human sample collection

This study included seven liver cancer patients from the University Medical Center in Mainz and the NIH Clinical Center. Detailed clinical information of this cohort can be found in Table S1. A total of 50 samples were collected from these patients. We conducted single-cell spatial transcriptome profiling of 15 samples. Additionally, we generated single-cell transcriptome data for the remaining samples.^20^ This study was approved by the ethics committee of the University Medical Center in Mainz and the National Institutes of Health.

### CosMx SMI Sample Preparation

Formalin-fixed, paraffin-embedded (FFPE) tissue sections were prepared for CosMx SMI profiling as described in He et al.^19^ Briefly, five-micron tissue sections were cut and placed on Leica BOND PLUS slides. Slides were baked overnight at 60 °C to improve tissue adherence to the slide. Tissues were deparaffinized in Xylene twice for 5 min each. Then, tissues were successively dehydrated in 100% ethanol twice for 5 min each and subsequently dried at 60 °C for 5 more min. Tissues were then prepared for in-situ hybridization (ISH) by heat-induced epitope retrieval (HIER) at 100 °C for 15 min using 1X Target retrieval solution (NanoString CMx Slide Prep Kit, FFPE RNA 121500006) in a Bio SB cooker. After target retrieval, tissue sections were rinsed with diethyl pyrocarbonate (DEPC)-treated water (DEPC H_2_O, ThermoFisher), washed in ethanol for 3 min and dried at room temperature for 30 min. Once slides dried, an incubation frame was placed over the tissue (CosMx FFPE Slide Prep Kit RNA). Tissue was then digested with Proteinase K (provided by NanoString) 3 μg /ml at 40 °C for 30 min. Tissue sections were rinsed twice with DEPC H_2_O, incubated in 1:400 diluted fiducials (Bangs Laboratories) in 2× saline sodium citrate and Tween (0.001% Tween 20, Teknova) for 5 min at room temperature and washed with 1× PBS (ThermoFisher) for 5 min. After digestion and fiducial placement, tissue sections were fixed with 10% neutral buffered formalin (NBF) for 1 min at room temperature. Fixed samples were rinsed twice with Tris-glycine buffer (0.1 M glycine, 0.1 M Tris-base in DEPC H_2_O) and once with 1X PBS for 5 min each before blocking with 100 mM *N*-succinimidyl (acetylthio) acetate (NHS-acetate, ThermoFisher) in NHS-acetate buffer (0.1 M NaP, 0.1% Tween PH 8 in DEPC H_2_O) for 15 min at room temperature. The sections were then rinsed with 2X saline sodium citrate (SSC) for 5 min. NanoString ISH probes were prepared by incubation at 95 °C for 2 min and placed on ice, and the ISH probe mix (1 nM 980 plex ISH probe, 10 nM Attenuation probes, 1X Buffer R, 0.1 U/μL SUPERase•In™ [Thermofisher] in DEPC H_2_O) was pipetted on the tissue slides. An adhesive SecureSeal Hybridization Chamber (Grace Bio-Labs) was placed over tissue. The hybridization chamber was sealed to prevent evaporation, and hybridization was performed at 37 °C for 16h-18h. Tissue sections were rinsed of excess probes in 2X SSCT for 1 min and washed twice in 50% formamide (VWR) in 2X SSC at 37 °C for 25 min, then twice with 2X SSC for 2 min at room temperature. Samples were then prepared for antibody cocktail incubation. First, slides were incubated with nuclear stain 1:40 (provided by NanoString) for 15 min at room temperature protected from light. After a short PBS wash, tissues were incubated with a 4-fluorophore-conjugated antibody cocktail against CD298/B2M (488 nm), PanCK (532 nm), CD45 (594 nm), and CD68 (647 nm) proteins and DAPI stain in the CosMx SMI instrument. Antibodies were also included in the CosMx slide preparation kit by NanoString. After one hour incubation at room temperature protected from light, slides were washed three times with 1X PBS 5 min each. A custom-made flow cell was affixed to the slide in preparation for loading onto the CosMx SMI instrument. Flow cells were then stored in SSC while the instrument is being prepared for loading the samples.

### CosMx imaging

RNA target readout on the CosMx SMI instrument was performed as described by He et al.^19^ Briefly, the assembled flow cell was loaded onto the instrument and Reporter Wash Buffer was flowed to remove air bubbles. A preview scan of the entire flow cell was taken, and fields of view (FOVs) were placed on the tissue to match regions of interest identified. In our case we selected as many FOVs needed to completely cover the whole tissue on the slide. RNA readout began by flowing 100 μl of Reporter Pool 1 into the flow cell and incubation for 15 min. Reporter Wash Buffer (1 mL) was flowed to wash unbound reporter probes, and Imaging Buffer was added to the flow cell for imaging. Eight Z-stack images (0.8 μm step size) for each FOV were acquired, and photocleavable linkers on the fluorophores of the reporter probes were released by UV illumination and washed with Strip Wash buffer. The fluidic and imaging procedure was repeated for the 16 reporter pools, and the 16 rounds of reporter hybridization-imaging were repeated multiple times to increase RNA detection sensitivity. Then, tissue and cell membrane morphology visualization was performed using oligonucleotide-conjugated antibodies hybridized to a specific pool of barcode reporters. After unbound antibodies and DAPI stain were washed with Reporter Wash Buffer, Imaging Buffer was added to the flow cell and eight Z-stack images for the 5 channels (4 antibodies and DAPI) were captured.

### Image processing

An in-build SMI data analysis pipeline was used to perform image processing and feature extraction^19^. In brief, the pipeline involves in three major steps: 1) the fiducials embedded in the samples were used to match with the fixed image reference to adjust for potential shifts; 2) the reporter signature locations were assigned to x, y, z axes along with the associated confidence; 3) the location information of individual transcripts was extracted and converted to a table. An in-build machine learning algorithm Cellpose was used to assign transcripts to cell locations.^44,45^ The transcript profiles of individual cells were generated based on the locations of the transcripts and the cell boundaries determined with Z-stack images of nuclear (DAPI) and CD298/B2M (surface) staining.

### CosMx™ SMI data quality-control and preprocessing

With the coordinates and the raw transcript counts of individual cells, we filtered cells based on the following criteria: ≥20 counts detected and ≥10 genes detected; <10% negative probes and ≤5 negative probe counts. Cells with outlier cell area (p-value<0.01) based on Grubb’s test were also removed for downstream analysis. A total of 2,347,589 cells passed the quality control with a median of 93 features and 174 counts per cell. Data normalization was performed using the R package Seurat (v5.1.0).

### Major cell type annotation

Major cell type annotation (T cells, B cells, myeloid cells, endothelial cells, fibroblasts, and epithelial cells [including hepatocytes, cholangiocytes, and tumor cells]) of the single-cell spatial data was based on label transfer using the scRNA-seq dataset.^46^ The R package Harmony (v1.2.0) was used to integrate the CosMx™ SMI data with the scRNA-seq data. The top 20 principal components (PCs) were used for the integration, and the top 20 harmony embeddings were used to cluster the cells in the integrated dataset with default parameters. For each derived cluster, the percentage of different cell types from the scRNA-seq dataset was calculated. Clusters were annotated based on the highest proportion of cell types from scRNA-seq. A cluster was considered as unclassified if the highest proportion < 50% or the ratio of the second-highest proportion against the highest proportion > 50%. The analyses were performed for each sample separately. The expression score for each cell type was calculated as the mean values of the cell type specific genes (Table S2).

Malignant cells were determined based on their transcriptomes and geographic locations in a tumor. For epithelial cells from each individual patient, we identified different clusters of epithelial cells using Seurat. Specifically, after data normalization, most variable features were selected based on mean expressions and standard variances of genes. Top 20 PCA components were used for detecting neighbors and louvain clusters of epithelial cells were determined using default parameters. Each cluster was subsequently mapped to its spatial location. Clusters of epithelial cells located within tumor regions based on the histological images were classified as malignant cells, while those in non-tumor regions were annotated as non-tumor epithelial cells.

### Consistency of CosMx™ SMI and scRNA-seq data

To compare the gene expression profiles between CosMx™ SMI and scRNA-seq data generated from the same patients, we grouped the cells into six major cell types including T cells, B cells, myeloid cells, endothelial cells, fibroblasts and epithelial cells. We selected the genes that were detected in both datasets and calculated the average expression of each gene in individual cell types. We further calculated the Pearson correlation coefficient (R function “cor.test”) between averaged gene expression profiles from CosMx™ SMI data and scRNA-seq data to measure the consistency of the two profiling approaches.

### Define tumor cell transcriptomic states based on gene modules

Due to the limited number of genes in the single-cell spatial data compared with scRNA-seq data, we determined gene modules in malignant cells based on scRNA-seq data and further applied those identified modules into the single-cell spatial data to define tumor cell transcriptomic states. Gene module identification was conducted based on a previous published non-negative matrix factorization (NMF)-based method.^5^ We included two liver cancer single-cell datasets to improve the robustness and sensitivity of gene module identification.^15,46^ For each sample, the top 500 highly variable genes (HVGs) were used for data scaling and the NMF analysis. The number of cells was down sampled to 1000 for the samples containing over 1000 tumor cells. Negative values were set to 0, and genes with expression values of 0 across all cells within each sample were discarded. We then performed NMF on the scaled gene expression matrix for each sample separately using the R package NMF (v0.27). The “nsNMF” method and “nndsvd” seeding method were applied. The rank was set to 10 if the number of malignant cells per sample was less than 400, and to 30 for samples containing 400 or more malignant cells. The contribution matrices of genes in different modules for each sample were obtained. A gene was assigned to a module if it showed the highest contribution to that module across all modules within a sample. Modules obtained from individual samples were further filtered by the Jaccard index. Gene modules with ≥ 5% overlap with at least two other modules were retained. For each paired gene, the number of individual tumor modules in which they co-existed (designated as the connection value) was calculated, and a gene-gene connection matrix was constructed for gene pairs. Genes with a connection value greater than three with at least four other genes were retained for downstream analysis. The graph was clustered using the infomap clustering method from the R package igraph (v2.0.3). The final modules with at least 5 genes were retained. Using this approach, we identified 11 modules across all samples. We annotated the gene modules based on gene ontology (GO) terms from MSigDB ^47^, tumor derived signatures from Neftel et al.,^48^ Puram et al.,^49^ Ji et al.,^50^ and Barkley et al.^5^ Hypergeometric distribution test was used to determine the enrichment of our identified consensus module genes in annotated gene sets with R function “phyper”. FDR method was used to adjust the p-values.

Based on the gene modules identified from scRNA-seq data, module scores of individual epithelial cells from CosMx™ SMI dataset were calculated using the method from Barkley et al.^5^ For each module, 10,000 randomized gene lists were generated, each containing the same number of genes as the module. The average gene expression for each random list and the module was then calculated. The score for the module was defined as:

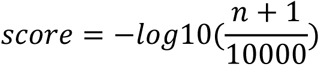

Where n represents the number of random gene set with higher average expression than the module. The score was further linearly rescaled to between 0 and 1. For the “Metabolism” module, only one gene was detected in the CosMx™ SMI dataset, so we excluded this module from further analysis. We removed cells with a score variance less than 0.15 and a mean score less than 0.2 across all modules to increase the confidence of module determination. The derived matrix was used to perform PCA and UMAP analyses. The epithelial cell clusters were determined by graph-based method using top 8 PCA components with default parameters in Seurat (v5.1.0). Average gene module scores were calculated for each cluster. Clusters with similar module scores were merged based on their hierarchical relationship and a total of 32 epithelial clusters were obtained. Epithelial clusters were annotated to specific transcriptomic states based on the highest gene module scores.

### Cell subtypes in non-malignant cells

For each non-malignant cell type (myeloid cells, T cells, B cells, endothelial cells, and fibroblasts) in the single-cell spatial transcriptome data, Harmony (v1.2.0) was used to integrate cells from different patients. Top 20 PCA components were used to perform harmony analysis and top 20 harmony embeddings were used for UMAP and clustering. Clusters were annotated based on differentially expressed genes. Marker genes from literature were used as reference in this analysis, including markers from Goveia et al.^51^ for endothelial subtypes, from Ma et al.^52^ for myeloid subtypes, from Cords et al.^53^ and Zhang et al.^54^ for fibroblast subtypes, from Morgan et al.^55^ for B cell subtypes, and from Guo et al.^56^, Zhang et al.^57^, Zheng et al.^58^, and Ma et al.^1^ for T cell subtypes.

### Align cell subtypes between CosMx™ SMI data and scRNA-seq data

To compare cell subtypes identified from the CosMx™ SMI data and the scRNA-seq data, we performed correlation analysis of the clusters identified from the two datasets. Specifically, we conducted cluster analysis of each non-epithelial cell type, i.e., myeloid cells, T cells, B cells, endothelial cells, and fibroblasts, from the scRNA-seq data with top 10 PCA and default resolutions. We selected the common genes from the CosMx™ SMI and scRNA-seq data to calculate the Pearson correlation coefficients between each pair of clusters from the two datasets. A CosMx™ SMI cluster was considered present in the scRNA-seq data if its correlation coefficient with any scRNA-seq cluster exceeded 0.5.

### Characterization of the surroundings of individual malignant cells

To characterize the surroundings of individual malignant cells, we determined the composition of surrounding cells for each targeted malignant cell in the CosMx™ SMI data. The surrounding cells are defined as those located within a 40 µm radius of the targeted cell. We included three layers of information to characterize the surrounding cell compositions, including major cell types, cell states/subtypes, and cell clusters. This vector of cell compositions was used as features to describe the targeted cell. We further performed PCA analysis of the surrounding composition matrix for all malignant cells and used the top 10 PCA components to generate a UMAP representation with default parameters. Clusters of malignant cells were determined using Louvain algorithm at a resolution of 0.6. Each cluster of malignant cells was annotated based on the composition of the surrounding cells.

To determine the distance for surrounding cell characterization, we tested the radii of 20 µm, 40 µm, 60 µm, 80 µm, and 100 µm. We performed pair-wise comparison of the derived results using a normalized mutual information (NMI) score. We also tested the significance of the comparisons based on a random shuffling strategy. We used 1000 times of randomly shuffled malignant cells as a background to calculate the p-value. Compared with 20 µm, malignant cell clusters are more stable with a larger radius. At a radius of 40 µm, the median number of surrounding cells for all malignant cells is 33. Considering that a larger radius will introduce a higher number of surrounding cells with a higher probability of noise to characterize a targeted cell, we selected a radius of 40 µm to determine surroundings of individual malignant cells in our liver cancer single-cell spatial data.

### Enrichment of tumor cell states on SDNs

In the CosMx™ SMI data, the enrichment of tumor cell states on SDNs was calculated as the proportions of each tumor cell state with a specific SDN. We used a random shuffling approach (1000 times of randomly shuffling of tumor cell states) to calculate p-values and to determine the significance of the enrichment. To identify significantly enriched tumor cell states, p-values were calculated in a stringent way as p-value= *n*/1000, where n represents number of cases with expected scores≥0.8*observed score. A p-value less than 0.05 was considered significant.

We used 39 samples from three publicly available 10X genomics Visium datasets from i.e., Liu et al.^32^, Wu et al.^6^, and Zhang et al.^33^ to validate the enrichment of tumor cell states on SDNs. Among them, 24 samples from the tumor core were included in our analysis. To determine tumor cell state for each spot, we generated reference pseudo-bulk gene signatures based on the CosMx™ SMI data. We averaged gene expression of the same tumor cell states to build an expression vector for each state. Genes with mean expression <0.1 or standard deviation <0.1 across all cell states were removed to retain the variable genes. We further selected the genes that were detected in the three datasets to ensure consistency. For each Visium dataset, Spearman correlations were then calculated between reference cell state signatures and the gene expression of each spot from the Visium data, resulting in a correlation matrix (Matrix I) of cell states and spots. High correlation values indicate a high probability of the spots containing tumor cells with the indicated cell states. To determine the SDNs for each spot in the 10X Visium datasets, we collected all cells related to each SDN in the CosMx data. The averaged gene expression related to each SDN was determined. Similar to the analysis of tumor cell states, we used the same criteria to filter genes and calculate correlations to determine the SDN of each spot. A correlation matrix (Matrix II) of SDNs and spots was generated. To infer the tumor cell state enrichment on SDNs, we calculated the Pearson correlation between tumor cell states (from Matrix I) and SDNs (from Matrix II) across all spots. The Pearson correlation coefficients were linearly rescaled to a range between 0 and 1.

### Physical distance between malignant cells to endothelial cells

For each malignant cell, the distance to the closest endothelial cell was calculated using R function “kNN” from dbscan (version 1.1-12). R function “kdtree” was used to search the nearest cell.

### Identification of tumor cell villages

To identify tumor cell villages, we constructed an undirected graph for tumor cells from each sample, where nodes represent tumor cells and edges indicate connections of tumor cells within 40 µm distance. The cell state and the SDN were represented by one-hot encoding and combined as node features. Graph Attention Network (GAT) model implemented in PyTorch (v1.21.1) was applied to train the connected graphs for node clustering. In contrast to other methods, GAT utilizes attention mechanisms. During training, the model learns the importance of each neighboring node’s features to the target node as attention coefficients for the target node. The coefficients are then normalized by SoftMax function to sum to 1. We used this attention scoring as a representation of interaction strength between nodes. These coefficients are used during feature aggregation in that the learned node feature is a weighted sum of neighbor information according to the attention coefficients. A 128-dimensional hidden representation was used for message passing between neighbors. Three convolutional layers was used in our analyses to aggregate neighborhood information of 3-hops. The Exponential Linear Unit (ELU) activation function was applied for each layer except for the last layer, followed by a 25% dropout rate for regularization. The model outputted a 15-dimension feature vector for each cell. To maximize the similarity of the embedded neighboring cells while minimizing that of distant cells, we used an evaluation method by Shiao et al.^53^ Specifically, a minibatch of 10,000 target cells was randomly sampled from the input graph. For each target cell, we sampled two types of cells: one directly connected to the target cell, and another chosen from a random position in the graph, with the latter assumed to be spatially unrelated to the target cell. The evaluation function maximizes similarity between neighboring cells and minimizes similarity of unrelated cells. The Adam optimization algorithm was used during training with a learning rate of 1e-5 for 100 epochs. We performed dimensional reduction on the derived embeddings from GAT for all the tumor cells using PCA. The top 10 PCA components were used for Louvain clustering at a resolution of 0.5, resulting in 10 tumor clusters. Clusters 9 and 10 were discarded from the downstream analysis due to a small number of cells, accounting for less than 1% of total malignant cells.

### Village graph of malignant cells

Village graph represents the compositions of tumor cell states and SDNs associated with each village. The sizes of the dots indicate the proportion of tumor cell states in each village. Tumor cell states with a proportion of at least 5% were included in the graph. Interaction weights between tumor cells were obtained from graph attention auto-encoder model described above, with higher values indicating stronger interactions. The interactions were visualized using the Fruchterman-Reingold layout. The outer ring stands for the SDNs, with different areas indicating the proportions of different SDNs related to each village.

### Define tumor villages using gene signatures

We determined gene signatures for each village based on differential gene expression analysis of the CosMx™ SMI data. Genes with fold changes ≥ 1 and p-values < 0.05 were selected as surrogate signatures for each village. To determine the specificity of the signatures to each village, we constructed a pseudo-bulk dataset based on the CosMx™ SMI data. For each tumor cell, surrounding cells within a 40 µm radius, measured by Euclidean distance, were collected. Average gene expression across cells was then calculated to generate pseudo-spot bulk gene expression. For each pseudo-spot, we determined the village scores as the mean expression of each village-related genes. Each pseudo-spot was assigned to the village with the highest village score. A confusion matrix was calculated between the predicted villages and ground truth from CosMx™ SMI data analysis. Accuracy, precision, recall and F1 score were calculated to measure the performance of our village detection model. F1 score was calculated as follows:

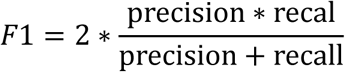

For the 10X Visium dataset, we used the same method to determine tumor cell villages for the spots. For each dataset, village scores were then linearly rescaled to a range between 0 and 1 across all spots. The confidence of village assignment for each spot was calculated as the difference between the highest village score and the second highest village score.

### Overall survival

For each bulk cohort, tumor samples were clustered based on the village scores. Hierarchical relationship of the samples was constructed using Pearson correlation as distance and “complete” agglomeration method. Samples in each cohort were grouped into three clusters based on the hierarchical relationship. Kaplan-Meier plots were generated and visualized using “Surv” function from R package Survival (v3.7-0).

### Cell-cell interactions using single-cell spatial transcriptomics data

To infer cell-cell communications from the CosMx™ SMI data, we used the COMMOT method^59^, with CellChatDB as the ligand-receptor (LR) database. For each sample, we included only LR pairs identified in ≥100 cells or in ≥5% of all tested cells. Cells within a 50 µm distance were considered for interaction calculations. To identify LR interactions associated with tumor cell states, we used a multi-step approach. First, LR scores were calculated for each tumor cell state using tl.cluster_communication function in COMMOT, with cells categorized into B cells, T cells, endothelial cells, epithelial cells, fibroblasts, myeloid cells, and 12 distinct tumor cell states. LR scores were considered significant if p-value < 0.05, determined by 1,000 cell label shuffles. LR scores were set to 0 for non-significant interactions and further averaged across samples to minimize sample bias. Next, LR scores between different tumor cell states were compared, pairs with a log2-fold change ≥ 0.5 and a signal in ≥ 10% of tumor cell states were selected. Finally, for each tumor cell state, a ranking score (score = P × FC) was calculated for each LR interaction based on the proportion of cells with the LR interaction (P) and the log2-fold change compared with other tumor cell states (FC), both rescaled between 0 and 1. The top three LR interactions for each tumor cell state were selected based on these ranking scores.

### Molecular co-dependencies in tumor cell villages

We developed a bioinformatic method to determine the molecular co-dependencies between tumor cells and their neighboring non-tumor cells (including CAFs, TAMs, TECs, T cells and B cells) within each tumor cell village. Specifically, for a given tumor-neighboring cell type pair in a tumor cell village, we identified the top 50 highly variable genes (HVGs) for each cell type. For each tumor cell–neighboring cell type pair, we randomly selected 1000 tumor cells, 1000 paired neighboring cells of a given cell type located based on a 40 µm distance. Pearson correlation coefficients were calculated between each pair of tumor-HVGs and non-tumor cell HVGs (each cell type separately). We repeated the random selections for 1000 times. The top 50 gene pairs with the highest correlations were selected for each tumor cell village. We further determined the correlations of these gene pairs within each tumor cell villages based on 40–80 µm, 80–160 µm, and 160–320 µm distance, and beyond each tumor cell village. The correlation scores were further linearly rescaled to a range between 0 to 1 within each village.

### Random forest model to assess molecular co-dependency

We used a random forest model to evaluate the importance of the identified gene pairs in each tumor cell village. A random forest model was built using expression profiles of the identified gene pairs. For each tumor cell, the expression of the paired genes in a corresponding neighboring cell type was averaged based on a 40 µm distance. We used two-thirds of the tumor cells in each village as the training set, with default parameters from the R package randomForest (v4.7-1.2). Tumor cell villages were used as classifier labels. The remaining one-third of the tumor cells served as the test set to evaluate model performance. To examine the importance of the gene pairs, we spatially shuffled the neighbors of tumor cells to construct perturbed neighboring expression profiles and assess the model’s performance on these perturbed profiles. Additionally, we spatially shuffled the neighbors of SPINK1^+^ tumor cells to specifically evaluate the impact of the SPINK1 gene pairs in Villages 3 and 6. The model-building and neighbor perturbation steps were repeated 100 times to mitigate potential random effects and ensure robustness of the results.

### Spatial co-localization of SPINK1^+^ spots and spots positive for given genes

For each sample in the 10X Visium dataset, we identified SPINK1^+^ spots as those with the top 25% SPINK1 expression across all spots. Genes of interest (GOI, e.g., BGN, COL1A1, COL1A2, COL4A1, COL5A1, IGFBP7, LUM, MGP, TPT1, and VIM)-positive spots were similarly defined using the top 25% expression threshold. For each SPINK1+ spot, we used the ‘kNN’ function from the R package dbscan (v1.2-0) to identify the nearest GOI-positive spots and calculate their distances. The distances for all SPINK1^+^ spots were averaged as the observed distance between SPINK1^+^ spots and GOI-positive spots for each sample. To assess significance, we generated 1000 random spot lists, each containing the same number of spots as GOI-positive spots. For each random list, ‘kNN’ was used to calculate the average distance of SPINK1+ spots and the random spots (random distance). P-values were calculated as p = n/1000, where n is the number of random distances equal to or greater than the observed distance for the given gene pair in a sample.

### Gene Set Enrichment Analysis (GSEA)

R package fgsea (v1.32.0) was used to perform GSEA analysis. Pathways with p-values <0.05 were considered as significantly enriched pathways.

### Cell-cell communications based on single-cell transcriptomics data

The function “nichenet_seuratobj_cluster_de” from the R package NicheNetR (v2.1.0) was used with default parameters to infer cell-cell communication in single-cell transcriptomics. The top 20 most significant ligands were selected to visualize the cell-cell communication between different cell groups.

### Data and code availability

The single-cell spatial transcriptome data generated from this study have been deposited in Zenodo (https://zenodo.org/doi/10.5281/zenodo.13773977). The processed scRNA-seq data of the patients in this study are available through the Gene Expression Omnibus (accession number <u>GSE189903</u>) and the raw sequencing data are available through dbGaP under accession number <u>phs003117.v1.p1</u>. The publicly available 10X genomics visium datasets used in this study include samples from Liu et al.^32^ (Mendeley Data: skrx2fz79n), Wu et al.^6^ (Genome Sequence Archive (GSA): HRA000437), and Zhang et al.^33^ (GEO accession: GSE238264). Other publicly available data include bulk transcriptomic data of <u>GSE14520</u> (LCI), GSE144269 (Mongolia), and the <u>TCGA</u> database (TCGA-LIHC). Code is available upon reasonable request.

