## supplementary files, figures and tables for "Tumor cell villages define the co-dependency of tumor and microenvironment in liver cancer"

**Figure S1. Quality control of the CosMx$^{TM}$ SMI data.**
(A) Boxplots of the number of genes or counts detected per cell in each sample (left two panels). Bar plots of the number of field of views (FOVs) and the number of cells in each sample (right panel).
(B) Expression of cell type-specific marker genes. Color indicates normalized expression and dot size indicates the fraction of cells expressing a certain gene.
(C) Protein intensities of Pan-CK (Pan-Cytokeratin), CD45, and CD68 in each cell type. P-values are calculated using Student's t-test. ****, p-value <0.001.
(D) Correlation of gene expression between CosMx$^{TM}$ SMI data and scRNA-seq data in individual cell types. Each dot represents a gene. Pearson correlation coefficients and p-values are shown for each cell type. ***, p-value < 0.001. Linear regression was applied.
(E) UMAP embeddings of malignant cells (left, colored by samples) and non-malignant cells (right, colored by cell types).

**Figure S2. Transcriptomic states of epithelial cells in liver cancer.**
(A) Heatmap of module-specific genes. Connections of genes were calculated as the number of individual tumor modules in which they co-occurred (see **Methods** for details).
(B) Recurrent genes modules, key representative genes, and abbreviations of the identified gene modules.
(C) UMAP embeddings of all epithelial cells (including malignant cells and non-malignant epithelial cells) from the CosMx$^{TM}$ SMI data based on gene module scores. Clusters were annotated based on module scores and marker genes.
(D) Proportion of each transcriptomic state in malignant cells (left) and non-malignant epithelial cells (right).
(E) Composition of malignant cell transcriptomic states in each tumor sample.

**Figure S3. Landscape of non-malignant cells.**
UMAP of each non-malignant cell type (left) and the expression of marker genes (right). In each UMAP, cell subtypes are indicated by colors. In the dot plots, color indicates normalized average gene expression, and the size of the dots stand for the fraction of cells expressing a certain gene. EC, endothelial cell PCV; post-capillary vein; LSEC, liver sinusoidal endothelial cell; CAF,

cancer-associated fibroblast; mCAF, matrix CAF; rCAF, reticular-like CAF; eCAF, EMT-like CAF; SMC, smooth muscle cell; tCAF, tumour-like CAF; ifnCAF, interferon-response CAF; iCAF, inflammatory CAF; apCAF, antigen-presenting CAF; TAM, tumor-associated macrophage; Prolif-TAM, proliferating TAM; TIM, tumor infiltrating monocyte; Reg-TAM, immune-regulatory TAM; Angio-TAM, pro-angiogenic TAM; INT monocyte, intermediate monocyte; LA-TAM, lipid-associated TAM; Inflam-TAM, inflammatory cytokine-enriched TAM; PC, plasma cell; Prolif-B, proliferative B cell; Teff, effector T cell; Tmem, Memory T cell; TEMRA, recently activated effector memory or effector T cell; Treg, regulatory T cell; Prolif-T, proliferative T.

**Figure S4. Comparison of cell clusters identified from the CosMx™ SMI data and the scRNA-seq data from the same set of liver cancer patients.**

Donut plots of cell subtypes and clusters identified from the CosMx™ SMI data. Cell subtypes or clusters found exclusively in the CosMx™ SMI data are shown in grey, while those identified in both CosMx™ SMI data and scRNA-seq data are shown in other colors. The proportion of cell subtypes or clusters in Fig. 2C from the CosMx™ SMI dataset is shown.

**Figure S5. SDNs of malignant cells.**

(A) Distribution of the number of surrounding cells for individual malignant cells with different radii.

(B) Normalized mutual information (NMI) score between malignant cell clusters with different radii. P-values were calculated based on 1,000 permutations. **, p-value <0.01.

(C) UMAP of malignant cell clusters based on the embeddings derived from graph attention networks in Figure 5B.

(D) Bubble plot of the proportions of each cell state (column) as the surroundings of each malignant cell cluster in (C). Each row represents the surroundings of one malignant cell cluster. Color and dot size indicate proportions.

**Figure S6. Crosstalk between malignant cell states and their local environments.**

Ligand-receptor interactions between each malignant cell state and their local environments.

**Figure S7. Validation of tumor cell villages in liver cancer.**

(A) Bubble plot of the compositions of different cells in each tumor village. Color and dot size indicate proportions.

(B) Proportions of tumor cell villages in each sample.

(C) Heatmap of the differentially expressed genes of each tumor village. Representative markers were indicated.

(D) Upset plot of the overlap of village-specific genes. Left: bar plot of the total number of genes (x axis) for each village (y axis). Right: intersection of genes among different villages. Each column represents a set of genes either unique to each village (dark dots) or shared between villages (connected dots). The number of genes in each set is indicated as the height of the bar with a number, while sets of shared genes are indicated using dots, with villages indicated on the left.

(E) Confusion matrix between the determined tumor villages using graph attention networks and those predicted based on village-specific marker genes.

(F) Performance of village prediction by village specific marker genes.

(G) A representative example of tumor villages in Liu et al. The alpha levels of the spot indicate the confidence of village prediction.

(H) Proportions of tumor villages in each tumor sample from three publicly available 10X Visium datasets.

(I) Hierarchical clustering of tumor samples in TCGA, Mongolia, and LCI cohorts based on village scores. Village score was determined as average expression of genes in (C).

**Figure S8. Spatial molecular co-dependencies in tumor cell villages.**

(A) Log2 foldchange (log2FC) of the expression of genes (row) in tumor cells compared with other cell types (column). Color indicates log2FC level. Right panel showing summed value of log2FC for each gene across columns.

(B) A chord plot of cell-cell communications between mCAFs (magenta)/other fibroblasts (blue) and SPINK1+ tumor cells (red), where line widths represent the interaction strength.

**Table S1.** Clinical and sample information.

**Table S2.** Genes for cell type annotation.

**Table S3.** Gene modules in epithelial cells.

**Table S4.** Gene signatures of each tumor cell village.

**Table S5.** Top gene pairs in each tumor cell village.

Figure S1

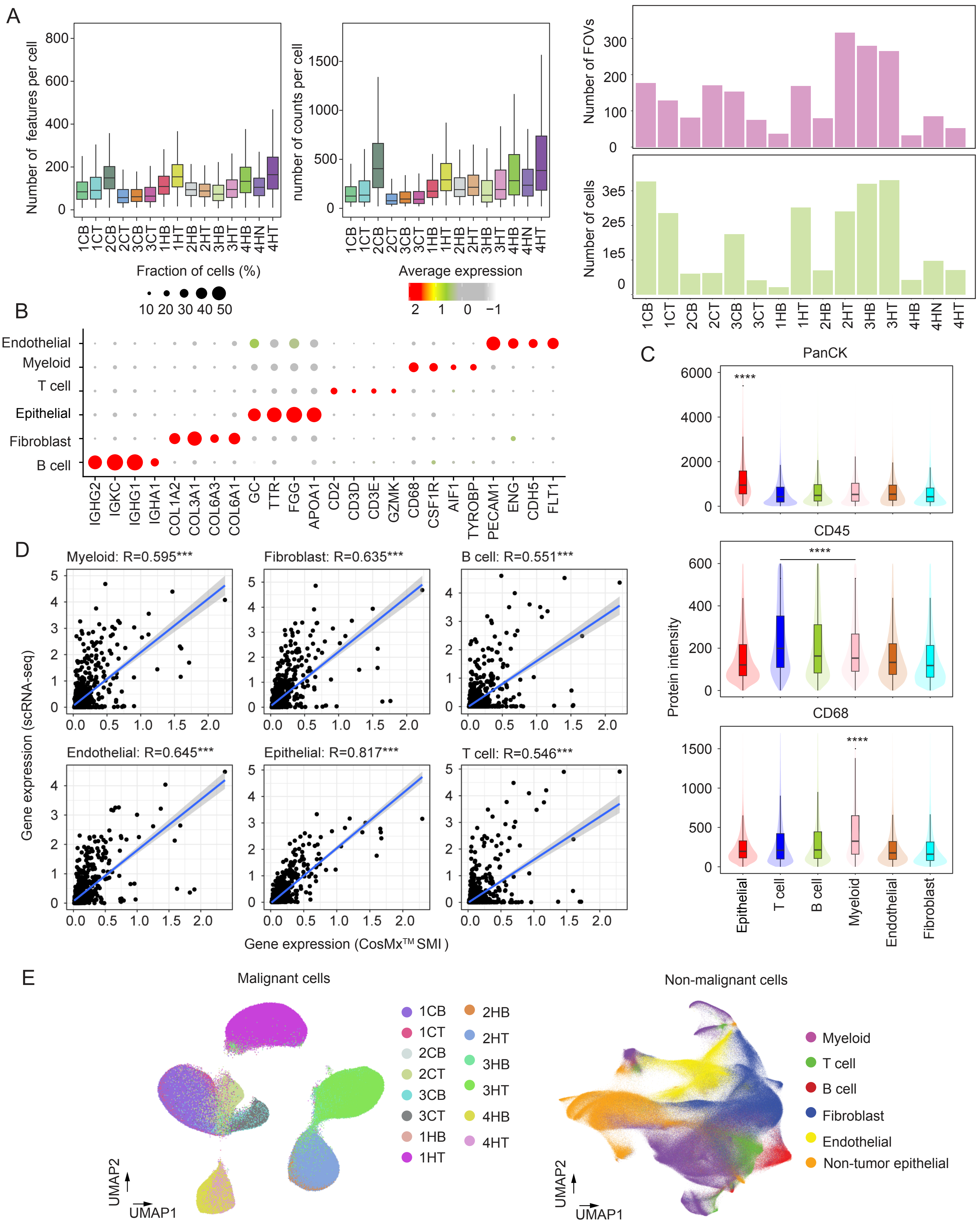
A
Number of features per cell
number of counts per cell
Number of FOVs
Number of cells
1CB 1CT 2CB 2CT 3CB 3CT 1HB 1HT 2HB 2HT 3HB 3HT 4HB 4HN 4HT
B
Fraction of cells (%)
10 20 30 40 50
Average expression
2 1 0 −1
Endothelial
Myeloid
T cell
Epithelial
Fibroblast
B cell
IGHG2 IGKC IGHG1 IGHA1 COL1A2 COL3A1 COL6A3 COL6A1 GC TTR FGG APOA1 CD2 CD3D CD3E GZMK CD68 CSF1R AIF1 TYROBP PECAM1 ENG CDH5 FLT1
C
PanCK
CD45
CD68
Protein intensity
Epithelial T cell B cell Myeloid Endothelial Fibroblast
D
Myeloid: R=0.595***
Fibroblast: R=0.635***
B cell: R=0.551***
Endothelial: R=0.645***
Epithelial: R=0.817***
T cell: R=0.546***
Gene expression (scRNA-seq)
Gene expression (CosMx™ SMI )
E
Malignant cells
1CB 1CT 2CB 2CT 3CB 3CT 1HB 1HT 2HB 2HT 3HB 3HT 4HB 4HT
UMAP2
UMAP1
Non-malignant cells
Myeloid
T cell
B cell
Fibroblast
Endothelial
Non-tumor epithelial

Figure S2

A

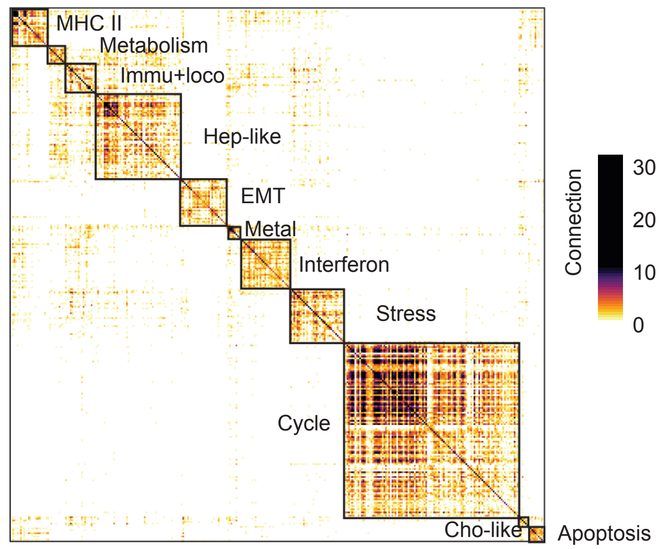


B

| Gene Modules | Signature genes | Abbreviation |
|---|---|---|
| Cell Cycle | HMGB2, TOP2A, CENPF | Cycle |
| Stress response | FOSB, JUN, HSPA6 | Stress |
| MHC II | IFI6, HLA-DRA, HLA-DRB | MHC II |
| Interferon response | STAT1, IFIT1, ISG20 | Interferon |
| EMT | VIM, COL4A1 | EMT |
| Metallothionein | MT2A, MT1G, MT1A | Metal |
| Immune response & locomotion | CCL20, CXCL5, ICAM1 | Immu+loco |
| Hepatocyte-like | ALB, SERPINA1, CYP2E1 | Hep-like |
| Cholangiocyte-like | KRT7, S100A6, S100A11 | Cho-like |
| Apoptosis | LTB, SLC2A3 | Apoptosis |
| Metabolic process | ALDH1A1, AKR1C1, GSTA2 | Metabolism |

C

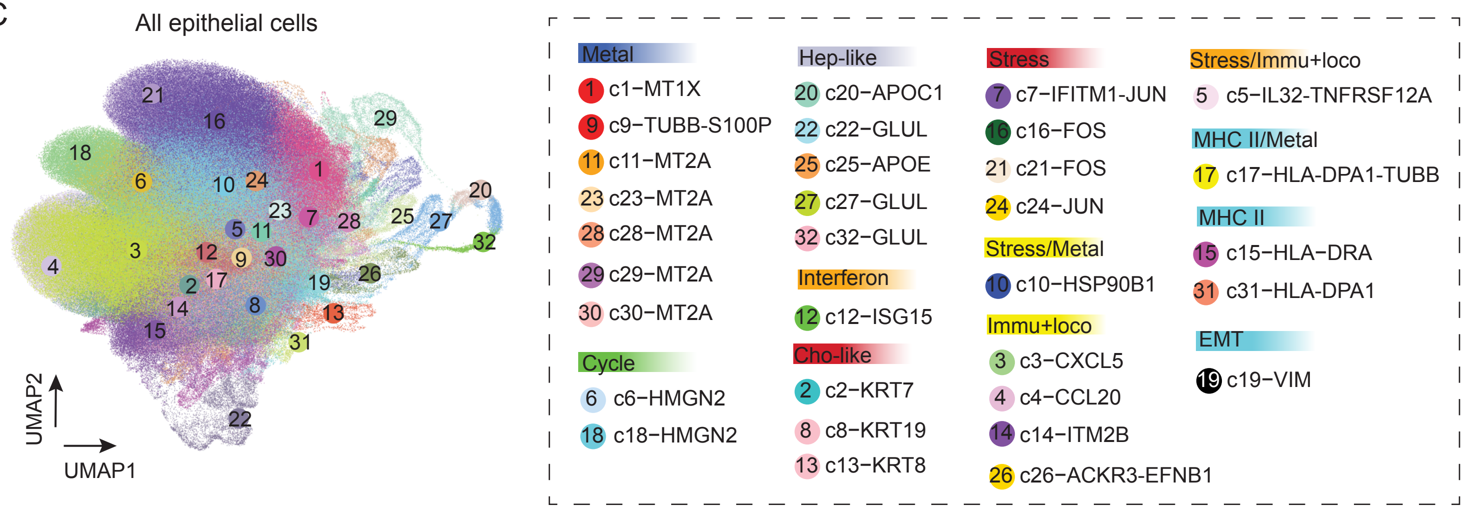


D

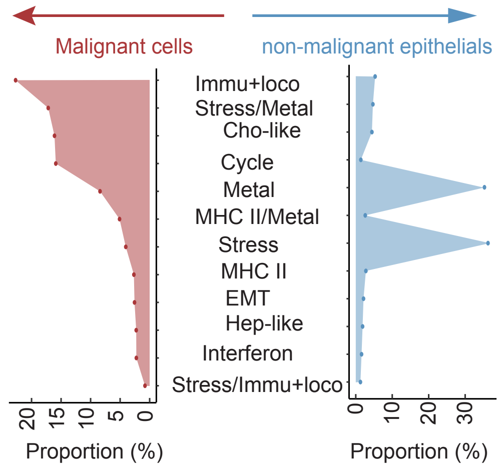


E

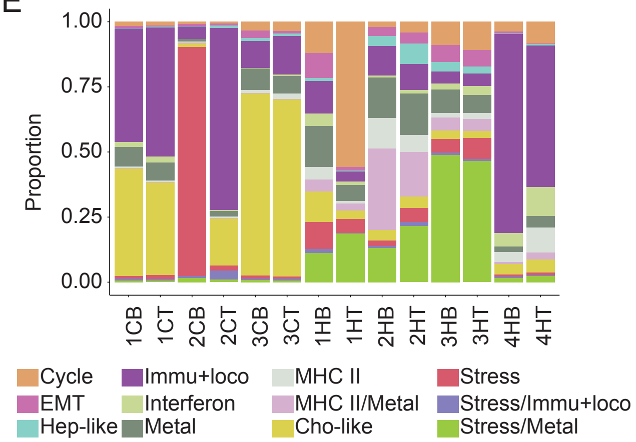

Figure S3

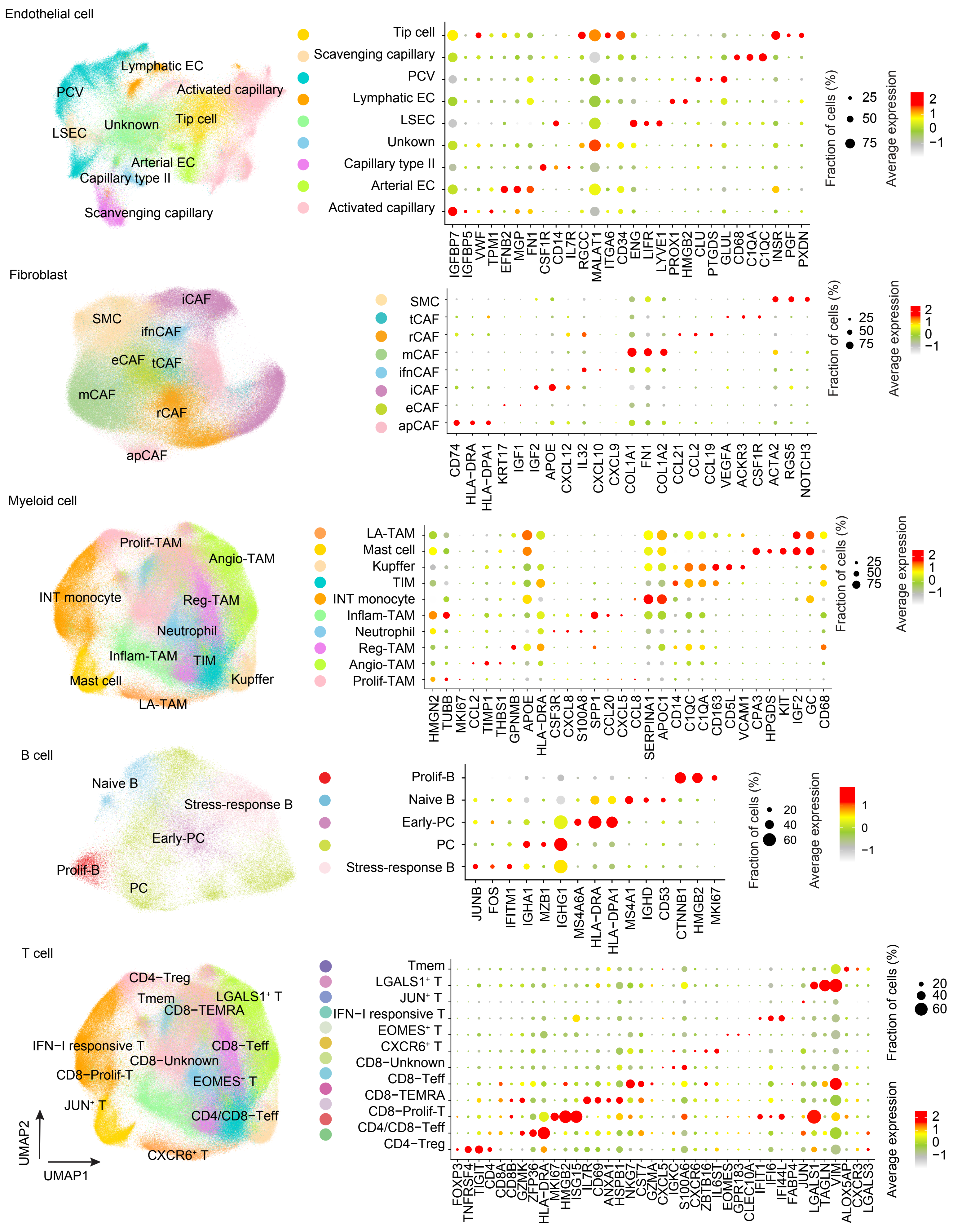
Endothelial cell
Lymphatic EC
PCV
Activated capillary
Unknown
Tip cell
LSEC
Arterial EC
Capillary type II
Scanvenging capillary
Tip cell
Scavenging capillary
PCV
Lymphatic EC
LSEC
Unkown
Capillary type II
Arterial EC
Activated capillary
IGFBP7 IGFBP5 VWF TPM1 EFNB2 MGP FN1 CSF1R CD14 IL7R RGCC MALAT1 ITGA6 CD34 ENG LIFR LYVE1 PROX1 HMGB2 CLU PTGDS GLUL CD68 C1QA C1QC INSR PGF PXDN
Fraction of cells (%)
25
50
75
Average expression
2
1
0
−1
Fibroblast
iCAF
SMC
ifnCAF
eCAF
tCAF
mCAF
rCAF
apCAF
SMC
tCAF
rCAF
mCAF
ifnCAF
iCAF
eCAF
apCAF
CD74 HLA−DRA HLA−DPA1 KRT17 IGF1 IGF2 APOE CXCL12 IL32 CXCL10 CXCL9 COL1A1 FN1 COL1A2 CCL21 CCL2 CCL19 VEGFA ACKR3 CSF1R ACTA2 RGS5 NOTCH3
Fraction of cells (%)
25
50
75
Average expression
2
1
0
−1
Myeloid cell
Prolif-TAM
Angio-TAM
INT monocyte
Reg-TAM
Neutrophil
Inflam-TAM
TIM
Kupffer
Mast cell
LA-TAM
LA-TAM
Mast cell
Kupffer
TIM
INT monocyte
Inflam-TAM
Neutrophil
Reg-TAM
Angio-TAM
Prolif-TAM
HMGN2 TUBB MKI67 CCL2 TIMP1 THBS1 GPNMB APOE HLA−DRA CSF3R CXCL8 S100A8 SPP1 CCL20 CXCL5 CCL8 SERPINA1 APOC1 CD14 C1QC C1QA CD163 CD5L VCAM1 CPA3 HPGDS KIT IGF2 GC CD68
Fraction of cells (%)
25
50
75
Average expression
2
1
0
−1
B cell
Naive B
Stress-response B
Early-PC
Prolif-B
PC
Prolif-B
Naive B
Early-PC
PC
Stress-response B
JUNB FOS IFITM1 IGHA1 MZB1 IGHG1 MS4A6A HLA−DRA HLA−DPA1 MS4A1 IGHD CD53 CTNNB1 HMGB2 MKI67
Fraction of cells (%)
20
40
60
Average expression
1
0
−1
T cell
CD4−Treg
Tmem
LGALS1⁺ T
CD8−TEMRA
IFN−I responsive T
CD8−Teff
CD8−Unknown
CD8−Prolif-T
EOMES⁺ T
JUN⁺ T
CD4/CD8−Teff
CXCR6⁺ T
UMAP2
UMAP1
Tmem
LGALS1⁺ T
JUN⁺ T
IFN−I responsive T
EOMES⁺ T
CXCR6⁺ T
CD8−Unknown
CD8−Teff
CD8−TEMRA
CD8−Prolif-T
CD4/CD8−Teff
CD4−Treg
FOXP3 TNFRSF4 TIGIT CD4 CD8A CD8B GZMK ZFP36 HLA−DRA MKI67 HMGB2 ISG15 IL7R CD69 ANXA1 HSPB1 NKG7 CST7 GZMA CXCL5 IGKC S100A6 CXCR6 ZBTB16 IL6ST EOMES GPR183 CLEC10A IFIT1 IFI6 IFI44L FABP4 JUN LGALS1 TAGLN VIM ALOX5AP CXCR3 LGALS3
Fraction of cells (%)
20
40
60
Average expression
2
1
0
−1

Figure S4

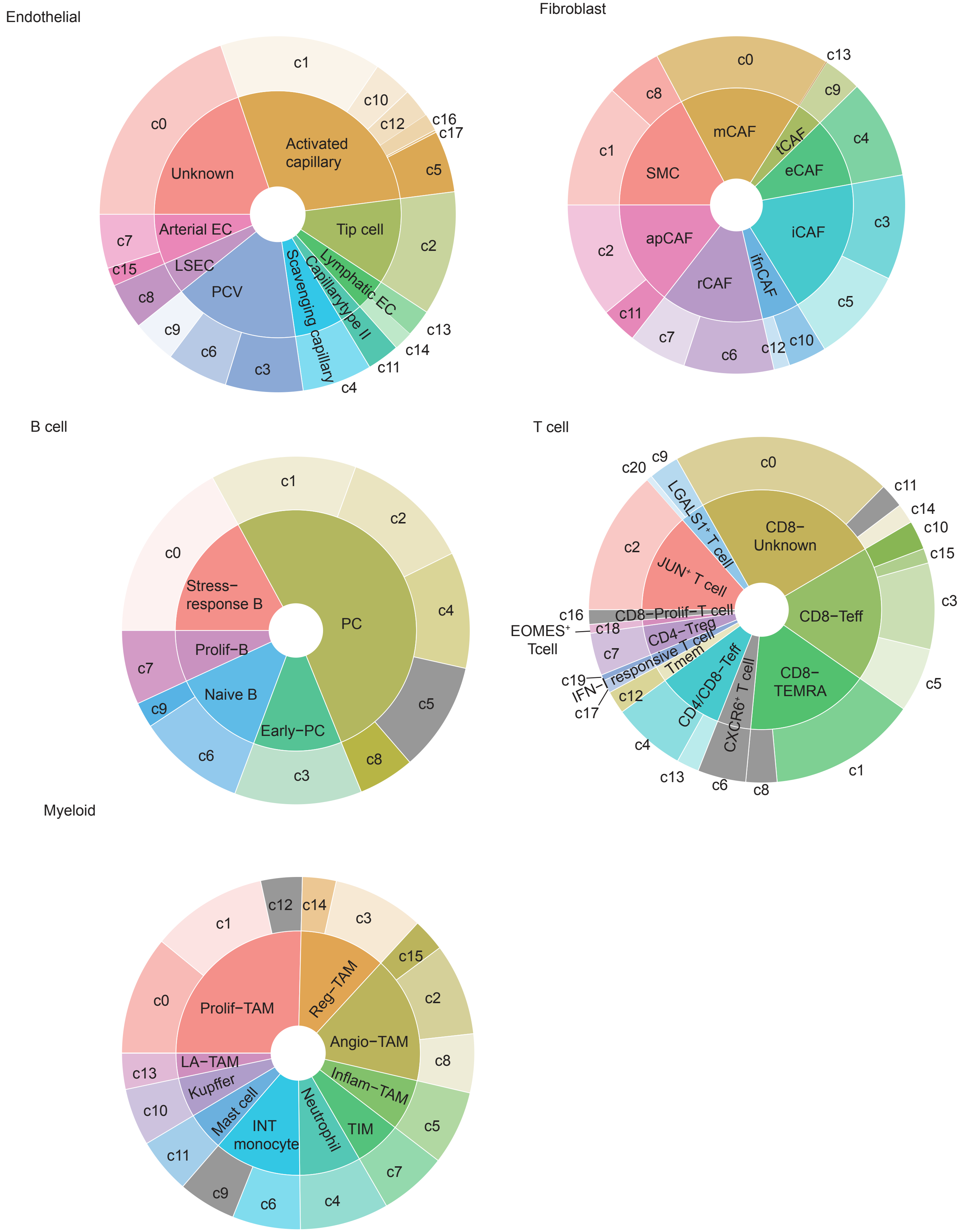

Endothelial
c1
c0
c10
c12
c16
c17
c5
Activated capillary
Unknown
Arterial EC
Tip cell
c2
c7
c15
LSEC
PCV
Scavenging capillary
Capillarytype II
Lymphatic EC
c8
c9
c6
c3
c4
c11
c14
c13
Fibroblast
c0
c13
c8
c9
mCAF
c1
SMC
tCAF
c4
eCAF
iCAF
c3
c2
apCAF
rCAF
ifnCAF
c5
c11
c7
c6
c12
c10
B cell
c1
c0
c2
Stress-response B
PC
c4
Prolif-B
c7
Naive B
Early-PC
c5
c9
c6
c3
c8
T cell
c20
c9
c0
c11
LGALS1⁺ T cell
CD8-Unknown
c14
c10
c15
c2
JUN⁺ T cell
c3
c16
CD8-Prolif-T cell
CD8-Teff
EOMES⁺ Tcell
c18
CD4-Treg
c7
IFN-I responsive T cell
Tmem
c19
c17
c12
CD4/CD8-Teff
CXCR6⁺ T cell
CD8-TEMRA
c5
c4
c13
c6
c8
c1
Myeloid
c12
c14
c3
c1
c15
Reg-TAM
c0
Prolif-TAM
c2
Angio-TAM
LA-TAM
c8
c13
Kupffer
Inflam-TAM
c10
Mast cell
INT monocyte
Neutrophil
TIM
c5
c11
c7
c9
c6
c4

Figure S5

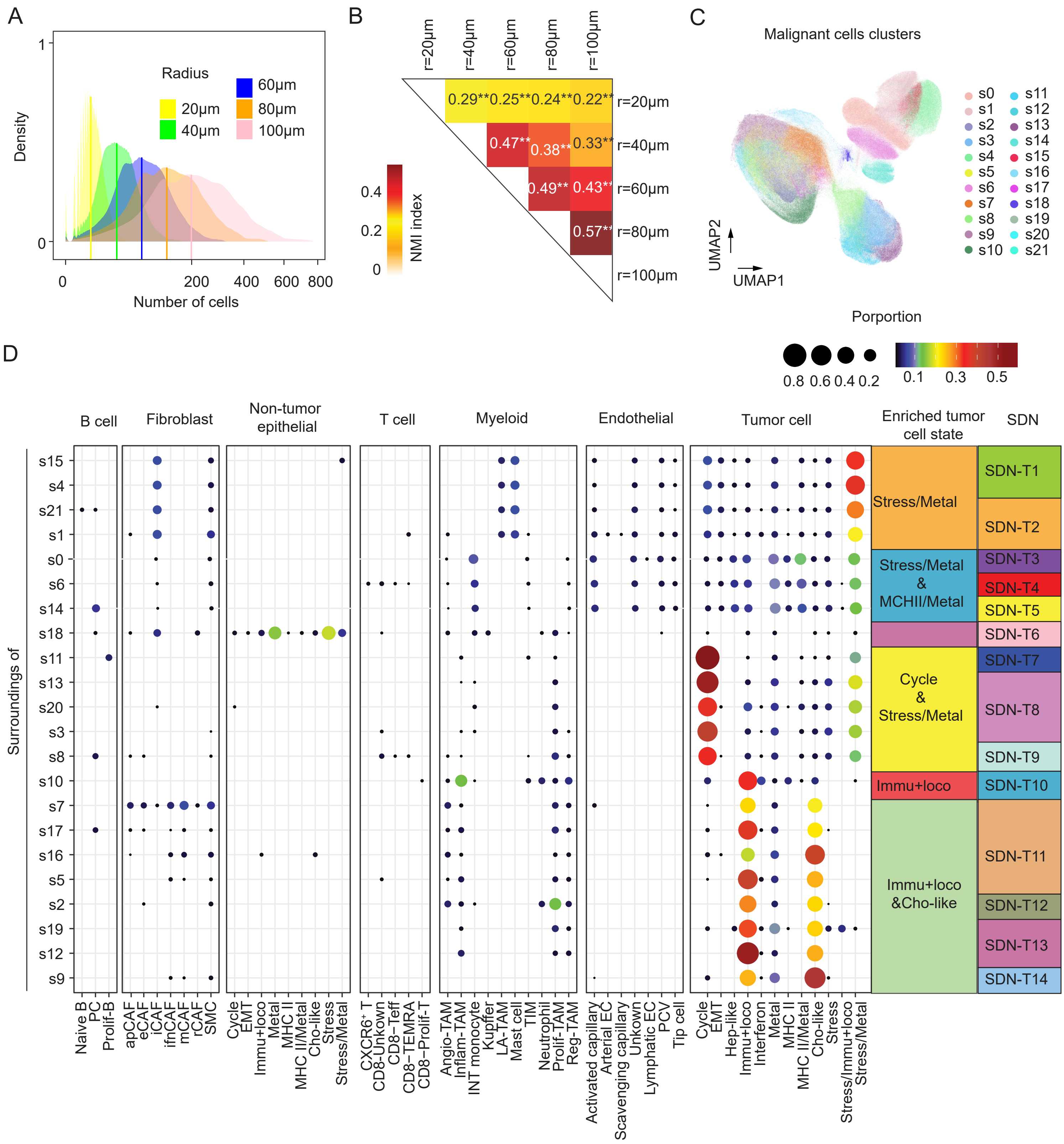
A
Radius
20µm
40µm
60µm
80µm
100µm
Density
Number of cells
B
r=20µm
r=40µm
r=60µm
r=80µm
r=100µm
0.29** 0.25** 0.24** 0.22**
0.47** 0.38** 0.33**
0.49** 0.43**
0.57**
NMI index
C
Malignant cells clusters
UMAP1
UMAP2
D
Porportion
B cell
Fibroblast
Non-tumor epithelial
T cell
Myeloid
Endothelial
Tumor cell
Enriched tumor cell state
SDN
Surroundings of
Stress/Metal
Stress/Metal & MCHII/Metal
Cycle & Stress/Metal
Immu+loco
Immu+loco &Cho-like
SDN-T1
SDN-T2
SDN-T3
SDN-T4
SDN-T5
SDN-T6
SDN-T7
SDN-T8
SDN-T9
SDN-T10
SDN-T11
SDN-T12
SDN-T13
SDN-T14
Naive B
PC
Prolif-B
apCAF
eCAF
iCAF
ifnCAF
mCAF
rCAF
SMC
Cycle
EMT
Immu+loco
Metal
MHC II
MHC II/Metal
Cho-like
Stress
Stress/Metal
CXCR6⁺ T
CD8-Unkown
CD8-Teff
CD8−TEMRA
CD8−Prolif-T
Angio-TAM
Inflam-TAM
INT monocyte
Kupffer
LA-TAM
Mast cell
TIM
Neutrophil
Prolif-TAM
Reg-TAM
Activated capillary
Arterial EC
Scavenging capillary
Unkown
Lymphatic EC
PCV
Tip cell
Hep-like
Interferon
Stress/Immu+loco

Figure S6

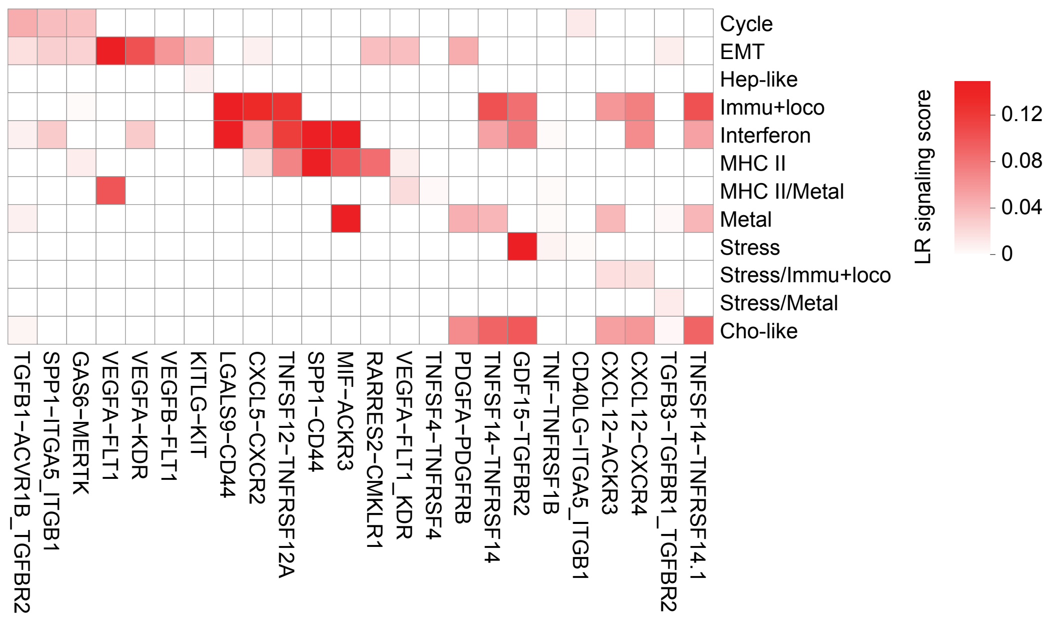
Cycle
EMT
Hep-like
Immu+loco
Interferon
MHC II
MHC II/Metal
Metal
Stress
Stress/Immu+loco
Stress/Metal
Cho-like
TGFB1−ACVR1B_TGFBR2
SPP1−ITGA5_ITGB1
GAS6−MERTK
VEGFA−FLT1
VEGFA−KDR
VEGFB−FLT1
KITLG−KIT
LGALS9−CD44
CXCL5−CXCR2
TNFSF12−TNFRSF12A
SPP1−CD44
MIF−ACKR3
RARRES2−CMKLR1
VEGFA−FLT1_KDR
TNFSF4−TNFRSF4
PDGFA−PDGFRB
TNFSF14−TNFRSF14
GDF15−TGFBR2
TNF−TNFRSF1B
CD40LG−ITGA5_ITGB1
CXCL12−ACKR3
CXCL12−CXCR4
TGFB3−TGFBR1_TGFBR2
TNFSF14−TNFRSF14.1
LR signaling score
0.12
0.08
0.04
0

### Figure S7

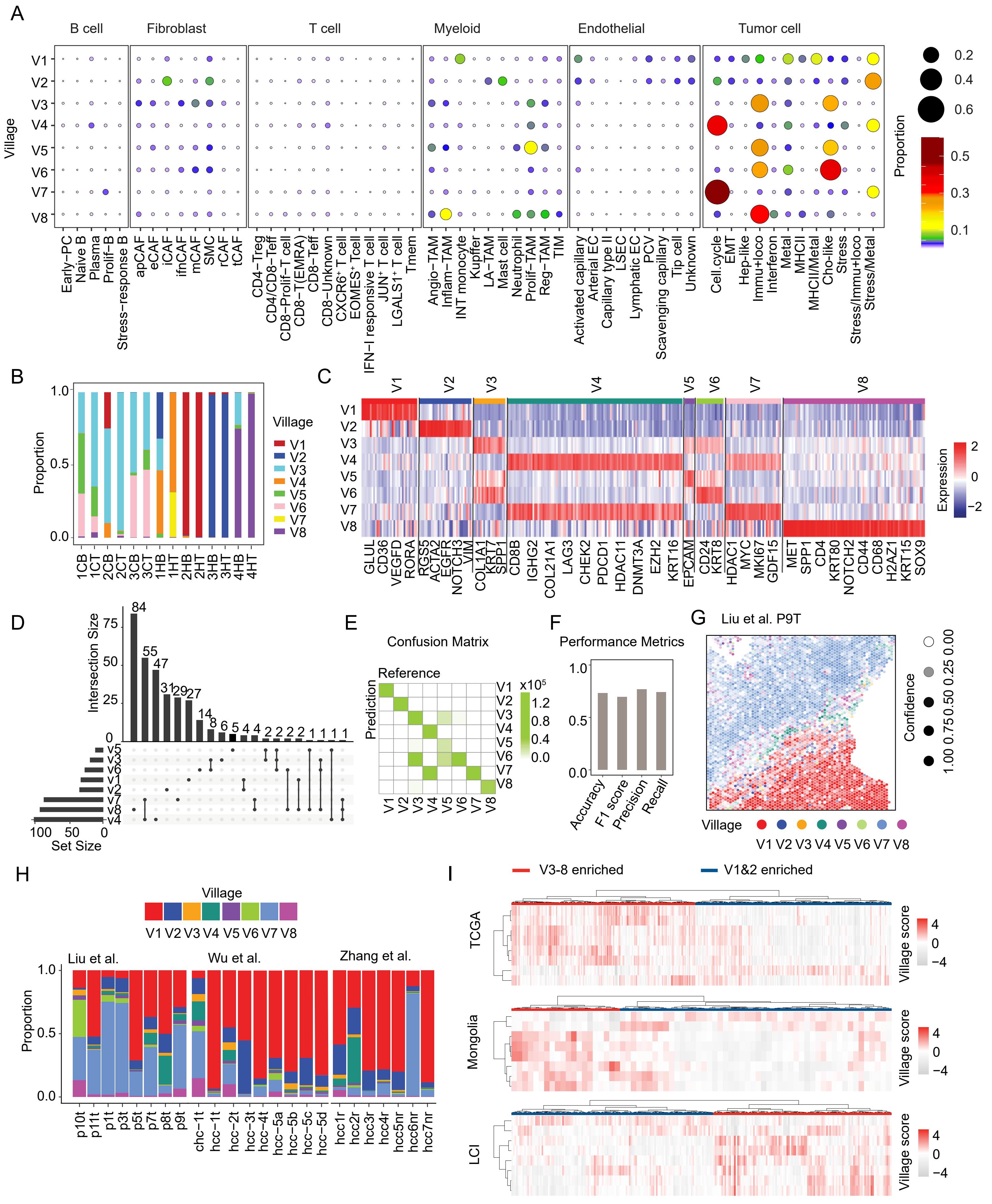
A
B cell
Fibroblast
T cell
Myeloid
Endothelial
Tumor cell
Village
V1
V2
V3
V4
V5
V6
V7
V8
Proportion
Early−PC
Naive B
Plasma
Prolif−B
Stress−response B
apCAF
eCAF
iCAF
ifnCAF
mCAF
SMC
rCAF
tCAF
CD4−Treg
CD4/CD8−Teff
CD8−Prolif−T cell
CD8−T(EMRA)
CD8−Teff
CD8−Unknown
CXCR6+ T cell
EOMES+ Tcell
IFN−I responsive T cell
JUN+ T cell
LGALS1+ T cell
Tmem
Angio−TAM
Inflam−TAM
INT monocyte
Kupffer
LA−TAM
Mast cell
Neutrophil
Prolif−TAM
Reg−TAM
TIM
Activated capillary
Arterial EC
Capillary type II
LSEC
Lymphatic EC
PCV
Scavenging capillary
Tip cell
Unknown
Cell.cycle
EMT
Hep−like
Immu+loco
Interferon
Metal
MHCII
MHCII/Metal
Cho−like
Stress
Stress/Immu+loco
Stress/Metal
B
Proportion
Village
1CB
1CT
2CB
2CT
3CB
3CT
1HB
1HT
2HB
2HT
3HB
3HT
4HB
4HT
C
Expression
GLUL
CD36
VEGFD
RORA
RGS5
ACTA2
EGFR
NOTCH3
VIM
COL1A1
KRT7
SPP1
CD8B
IGHG2
COL21A1
LAG3
CHEK2
PDCD1
HDAC11
DNMT3A
EZH2
KRT16
EPCAM
CD24
KRT8
HDAC1
MYC
MKI67
GDF15
MET
SPP1
CD4
KRT80
NOTCH2
CD44
CD68
H2AZ1
KRT15
SOX9
D
Intersection Size
Set Size
E
Confusion Matrix
Reference
Prediction
x10^5
F
Performance Metrics
Accuracy
F1 score
Precision
Recall
G
Liu et al. P9T
Confidence
H
Village
Proportion
Liu et al.
Wu et al.
Zhang et al.
p10t
p11t
p1t
p3t
p5t
p7t
p8t
p9t
chc−1t
hcc−1t
hcc−2t
hcc−3t
hcc−4t
hcc−5a
hcc−5b
hcc−5c
hcc−5d
hcc1r
hcc2r
hcc3r
hcc4r
hcc5nr
hcc6nr
hcc7nr
I
V3-8 enriched
V1&2 enriched
TCGA
Mongolia
LCI
Village score

Figure S8

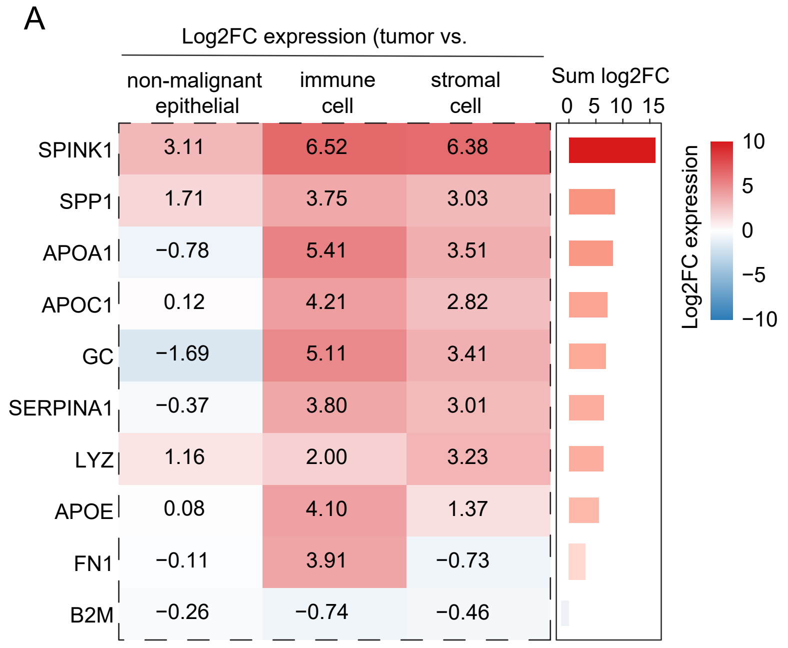
A
Log2FC expression (tumor vs.
non-malignant epithelial
immune cell
stromal cell
Sum log2FC
0 5 10 15
SPINK1 3.11 6.52 6.38
SPP1 1.71 3.75 3.03
APOA1 −0.78 5.41 3.51
APOC1 0.12 4.21 2.82
GC −1.69 5.11 3.41
SERPINA1 −0.37 3.80 3.01
LYZ 1.16 2.00 3.23
APOE 0.08 4.10 1.37
FN1 −0.11 3.91 −0.73
B2M −0.26 −0.74 −0.46
Log2FC expression
10
5
0
−5
−10

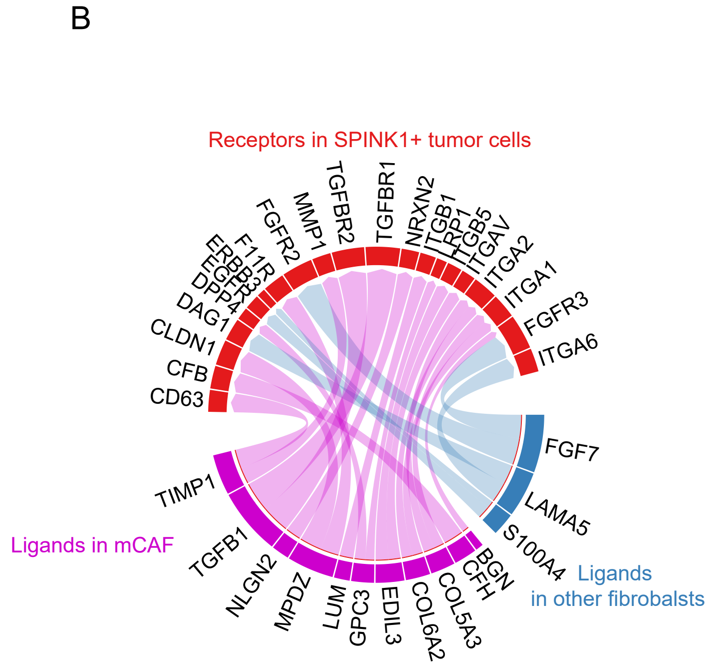
B
Receptors in SPINK1+ tumor cells
MMP1
TGFBR2
TGFBR1
NRXN2
ITGAV
ITGA2
ITGA1
FGFR3
ITGA6
FGFR2
DAG1
CLDN1
CFB
CD63
FGF7
LAMA5
S100A4
Ligands in other fibrobalsts
Ligands in mCAF
TIMP1
TGFB1
NLGN2
MPDZ
LUM
GPC3
EDIL3
COL6A2
COL5A3
BGN
CFH

Table S1. Clinical and sample information.

| Patient ID | Sample ID | CosMx | scRNA-seq | Age (years)[a] | Gender | Etiology | Diagnosis | Stage | Tumor size[b] | Vascular invasion | AFP (ng/ml) | CA19-9 (U/ml) |
|---|---|---|---|---|---|---|---|---|---|---|---|---|
| 1C | T1 | | Y | 75 | female | none | iCCA | II | 7.5 | yes | - | 89 |
| | T2 | | Y | | | | | | | | | |
| | T3 | | Y | | | | | | | | | |
| | T4 | Y | | | | | | | | | | |
| | B1 | | Y | | | | | | | | | |
| | B2 | Y | | | | | | | | | | |
| | N1 | | Y | | | | | | | | | |
| 2C | T1 | | Y | 71 | female | MASH | iCCA | I | 7.5 | no | <2.0 | 2.9 |
| | T2 | | Y | | | | | | | | | |
| | T3 | | Y | | | | | | | | | |
| | T4 | Y | | | | | | | | | | |
| | B1 | | Y | | | | | | | | | |
| | B2 | Y | | | | | | | | | | |
| | N1 | | Y | | | | | | | | | |
| 3C | T1 | | Y | 72 | male | MASH | iCCA | IVA | 5.5 | no | 6.3 | 3901 |
| | T2 | | Y | | | | | | | | | |
| | T3 | | Y | | | | | | | | | |
| | T4 | Y | | | | | | | | | | |
| | B1 | | Y | | | | | | | | | |
| | B2 | Y | | | | | | | | | | |
| | N1 | | Y | | | | | | | | | |
| 1H | T1 | | Y | 70 | male | none | HCC | II | 5.5 | yes | 323 | 2.5 |
| | T2 | | Y | | | | | | | | | |
| | T3 | | Y | | | | | | | | | |
| | T4 | Y | | | | | | | | | | |
| | B1 | | Y | | | | | | | | | |
| | B2 | Y | | | | | | | | | | |
| | N1 | | Y | | | | | | | | | |
| 2H | T1 | | Y | 77 | female | MAFLD | HCC | I | 7.7 | no | <2.0 | - |
| | T2 | | Y | | | | | | | | | |
| | T3 | | Y | | | | | | | | | |
| | T4 | Y | | | | | | | | | | |
| | B1 | | Y | | | | | | | | | |
| | B2 | Y | | | | | | | | | | |
| | N1 | | Y | | | | | | | | | |
| 3H | T1 | | Y | 63 | male | MASH | HCC | II | 24.5 | yes | 240144 | 2.1 |
| | T2 | | Y | | | | | | | | | |
| | T3 | | Y | | | | | | | | | |
| | T4 | Y | | | | | | | | | | |
| | B1 | | Y | | | | | | | | | |
| | B2 | Y | | | | | | | | | | |
| | N1 | | Y | | | | | | | | | |
| 4H | T1 | | Y | 76 | male | HCV,HIV | HCC | I | 3 | no | 16.1 | 0.8 |
| | T2 | | Y | | | | | | | | | |
| | T3 | | Y | | | | | | | | | |
| | T4 | Y | | | | | | | | | | |
| | B1 | | Y | | | | | | | | | |
| | B2 | Y | | | | | | | | | | |
| | N1 | | Y | | | | | | | | | |
| | N2 | Y | | | | | | | | | | |

[a]Age at time of tissue collection used for this study.
[b]Tumor size determined by radiologic measurements.

Table S2. Gene markers of cell types

| Myeloid | T cell | B cell | Fibroblast | Endothelial | Epithelial |
|---|---|---|---|---|---|
| CD68 | CD2 | CD79A | COL1A2 | PECAM1 | EPCAM |
| CTSG | CD3E | IGHM | FAP | VWF | KRT14 |
| ELANE | CD3D | P2RX5 | DCN | ENG | KRT17 |
| MPO | CD3G | CD19 | COL3A1 | CDH5 | KRT5 |
| PRTN3 | CD8A | MS4A1 | COL6A1 | FABP5 | KRT19 |
| CSF3R | FOXP3 | IGHA1 | COL6A3 | ESAM | KRT8 |
| CXCL8 | CD8B | IGHG2 | COL1A1 | FLT1 | KRT16 |
| S100A8 | CD40LG | MZB1 | THBS2 | | KRT18 |
| | ICOS | JCHAIN | FN1 | | KRT15 |
| | CTLA4 | CD2 | CDH11 | | SFN |
| | IL2RA | CD22 | | | KRT7 |
| | CD4 | CD40 | | | KRT20 |
| | NOTCH3 | CD69 | | | FGG |
| | IFIT3 | CD70 | | | SCGB3A1 |
| | LTB | CD80 | | | CEACAM1 |
| | CXCR6 | CD86 | | | ST6GAL1 |
| | IL7R | TNFRSF9 | | | ITGB4 |
| | ITK | TNFSF4 | | | IL1R1 |
| | PTPRCAP | TNFRSF13B | | | CDH1 |
| | RORA | PDCD1 | | | KRT1 |
| | CD69 | IGHD | | | ICAM1 |
| | CXCR4 | LTB | | | ITGAL |
| | BCL2 | CD14 | | | CD2 |
| | CTSW | LAIR1 | | | ITGA5 |
| | TXK | IFIT3 | | | ITGA2 |
| | LAG3 | CD27 | | | ITGA1 |
| | GZMK | IRF4 | | | FZD6 |
| | GZMB | LY6D | | | AGR2 |
| | CD28 | VPREB3 | | | APOA1 |
| | KLRB1 | TNFRSF17 | | | BRCA1 |
| | CCR2 | FCRLA | | | KRT13 |
| | IL2RB | FKBP11 | | | LTF |
| | IL18R1 | CXCR4 | | | PSCA |
| | TNFRSF4 | BIRC3 | | | TTR |
| | CCL20 | IGKC | | | PIGR |
| | CLEC2D | BST1 | | | PGR |
| | S100A4 | IGHG1 | | | MECOM |
| | CCL5 | CD38 | | | CXCL10 |
| | CD81 | PTPRC | | | CCL20 |
| | TCF7 | CCR7 | | | CXCL17 |
| | CCR7 | CD55 | | | AQP3 |
| | GZMA | CD74 | | | |
| | JUNB | CD52 | | | |
| | DUSP2 | | | | |
| | IFNG | | | | |
| | CD52 | | | | |
| | BRAF | | | | |

Table S3. Genes in different modules

| MHC II | Metabolism | Immu+loco | Hep-like | EMT | Metal | Interferon | Stress | Cycle | Cho-like | Apoptosis |
|---|---|---|---|---|---|---|---|---|---|---|
| CD74 | ADH1C | LCN2 | APCS | ID3 | S100P | GBP1 | ATF3 | FAM111B | S100A6 | CCL5 |
| HLA-DRA | AKR1B10 | TSPAN8 | CRP | VIM | MT1X | STAT1 | GADD45B | CENPU | S100A11 | CD52 |
| HLA-DRB1 | PTGR1 | C15orf48 | C4BPA | TUBA1A | MT2A | GBP4 | CXCL2 | TUBA1B | LGALS1 | S100A4 |
| HLA-DPB1 | GSTA1 | CCL20 | APOB | TIMP3 | MT1E | CTSS | NFKBIA | TYMS | EMP3 | LTB |
| IFI6 | ALDH1A1 | CXCL8 | ITIH3 | FLNA | MT1G | CXCL9 | JUN | CDCA8 | PHLDA2 | SRGN |
| ARHGDIB | AKR1C2 | CXCL5 | FGB | IGFBP7 | MT1F | CXCL10 | HSPA6 | CDC20 | KRT7 | CORO1A |
| PSMB9 | SPP1 | ICAM1 | FGA | TAGLN | TUBB | IRF1 | HES1 | DEPDC1 | | RGCC |
| HLA-DQB1 | AKR1C1 | CXCL6 | FGG | RBP7 | MT1A | UBD | FST | ATAD2 | | KLRB1 |
| HLA-DPA1 | GSTA2 | BIRC3 | ORM1 | SPARCL1 | MT1M | TAP1 | IER3 | PSRC1 | | HCST |
| LYZ | ALDOB | SOD2 | ORM2 | AQP1 | | ISG15 | PCK1 | NUF2 | | SLC2A3 |
| FCER1G | | TM4SF1 | HPX | GNG11 | | MX1 | GDF15 | CKS1B | | |
| HLA-DQA1 | | CXCL1 | NNMT | ENG | | IFIT2 | CXCL3 | ASPM | | |
| HLA-DRB5 | | IL32 | HP | VWF | | IFIT3 | PDK4 | CENPF | | |
| C1QA | | PLA2G2A | LBP | CDH5 | | IFITM1 | TRIB1 | CENPA | | |
| LST1 | | SAA2 | APOE | RAMP2 | | RARRES3 | DUSP1 | CKAP2L | | |
| AIF1 | | SAA1 | SLC2A2 | CLEC14A | | ISG20 | NR4A1 | HJURP | | |
| TYROBP | | TNFRSF12A | CHI3L1 | PECAM1 | | LAP3 | FOS | SGO1 | | |
| CCL3 | | GPX2 | HRG | TCF4 | | SAMD9L | RASD1 | TACC3 | | |
| C1QC | | MMP7 | GC | RGS5 | | IFIT1 | CYR61 | CENPE | | |
| C1QB | | | IGFBP1 | SPARC | | IFI44L | JUNB | CCNA2 | | |
| HLA-DMA | | | APOC1 | ACTA2 | | CX3CL1 | EGR1 | HMGB2 | | |
| MS4A7 | | | TF | MGP | | TRIM31 | KLF6 | CCNB1 | | |
| LAPTM5 | | | APOM | COL4A1 | | EPSTI1 | IER2 | PTTG1 | | |
| | | | ADH1B | TMSB4X | | CMPK2 | CTGF | CENPW | | |
| | | | APOC3 | PLVAP | | CXCL11 | HSPA1A | KIF4A | | |
| | | | APOA1 | COL4A2 | | ETV7 | HSPA1B | CKS2 | | |
| | | | AMBP | KDR | | UBE2L6 | FAM46A | CDK1 | | |
| | | | ITIH2 | IGFBP5 | | SAMD9 | BCO2 | CEP55 | | |
| | | | RBP4 | | | OAS3 | FOSB | MKI67 | | |
| | | | LRG1 | | | WARS | DNAJB1 | FOXM1 | | |
| | | | TIMP1 | | | IFI35 | LDLR | CDCA3 | | |
| | | | FABP1 | | | | RHOB | TROAP | | |
| | | | ALB | | | | WEE1 | CDKN3 | | |
| | | | REG1A | | | | | KNSTRN | | |
| | | | FGL1 | | | | | NUSAP1 | | |
| | | | SERPINA1 | | | | | CCNB2 | | |
| | | | AHSG | | | | | PLK1 | | |
| | | | TTR | | | | | CENPN | | |
| | | | CP | | | | | AURKB | | |
| | | | C9 | | | | | TOP2A | | |
| | | | LEAP2 | | | | | KPNA2 | | |
| | | | PGC | | | | | TK1 | | |
| | | | A2M | | | | | BIRC5 | | |
| | | | VTN | | | | | TPX2 | | |
| | | | CYP2E1 | | | | | MYBL2 | | |
| | | | APOA2 | | | | | UBE2C | | |
| | | | A1BG | | | | | UBE2S | | |
| | | | INSIG1 | | | | | RRM2 | | |
| | | | SERPIND1 | | | | | SMC4 | | |
| | | | HPR | | | | | GTSE1 | | |
| | | | NAT8 | | | | | HIST1H4C | | |
| | | | CFHR1 | | | | | ANLN | | |
| | | | UGT2B4 | | | | | MCM4 | | |
| | | | | | | | | E2F8 | | |
| | | | | | | | | HELLS | | |
| | | | | | | | | PKMYT1 | | |
| | | | | | | | | CENPM | | |
| | | | | | | | | SPC25 | | |

| MHC II | Metabolism | Immu+loco | Hep-like | EMT | Metal | Interferon | Stress | Cycle | Cho-like | Apoptosis |
|---|---|---|---|---|---|---|---|---|---|---|
| | | | | | | | | DTYMK | | |
| | | | | | | | | MAD2L1 | | |
| | | | | | | | | KIF20A | | |
| | | | | | | | | KIFC1 | | |
| | | | | | | | | ZWINT | | |
| | | | | | | | | OIP5 | | |
| | | | | | | | | MELK | | |
| | | | | | | | | PRC1 | | |
| | | | | | | | | PRR11 | | |
| | | | | | | | | TUBA1C | | |
| | | | | | | | | NDC80 | | |
| | | | | | | | | AURKA | | |
| | | | | | | | | KIF2C | | |
| | | | | | | | | CDT1 | | |
| | | | | | | | | KIF14 | | |
| | | | | | | | | UBE2T | | |
| | | | | | | | | NEK2 | | |
| | | | | | | | | ECT2 | | |
| | | | | | | | | TUBB4B | | |
| | | | | | | | | CLSPN | | |
| | | | | | | | | MCM5 | | |
| | | | | | | | | DTL | | |
| | | | | | | | | MCM7 | | |
| | | | | | | | | HMMR | | |
| | | | | | | | | DHFR | | |
| | | | | | | | | GINS2 | | |
| | | | | | | | | H2AFZ | | |
| | | | | | | | | RAD51AP1 | | |
| | | | | | | | | STMN1 | | |
| | | | | | | | | DLGAP5 | | |
| | | | | | | | | KIF23 | | |
| | | | | | | | | NCAPG | | |
| | | | | | | | | PBK | | |
| | | | | | | | | CKAP2 | | |
| | | | | | | | | PHF19 | | |
| | | | | | | | | KIF20B | | |
| | | | | | | | | UHRF1 | | |
| | | | | | | | | CENPK | | |
| | | | | | | | | MXD3 | | |
| | | | | | | | | ARL6IP1 | | |
| | | | | | | | | FAM83D | | |
| | | | | | | | | TCF19 | | |
| | | | | | | | | HMGN2 | | |
| | | | | | | | | H2AFX | | |
| | | | | | | | | BUB1 | | |
| | | | | | | | | MND1 | | |
| | | | | | | | | RACGAP1 | | |
| | | | | | | | | ARHGAP11A | | |
| | | | | | | | | KIAA0101 | | |
| | | | | | | | | FAM64A | | |
| | | | | | | | | SGOL1 | | |

Table S4. Gene markers of villages

| village_1 | village_2 | village_3 | village_4 | village_5 | village_6 | village_7 | village_8 |
|---|---|---|---|---|---|---|---|
| TTR | IGF2 | OLFM4 | CD8B | CXCL5 | KRT19 | STMN1 | MET |
| SPARCL1 | PTGDS | LUM | NRXN3 | REG1A | SPINK1 | LGR5 | FCGBP |
| CHI3L1 | GC | CRP | IGHG2 | CRP | S100A6 | COL21A1 | S100A8 |
| SELENOP | FGG | DCN | TOX | BIRC3 | KRT23 | CD8B | PHLDA2 |
| GLUL | BASP1 | MMP7 | COL21A1 | S100A6 | IL18 | ENO1 | CXCL10 |
| GC | ARG1 | S100A6 | BMP7 | CXCL8 | CEACAM6 | G0S2 | TYROBP |
| IL13RA1 | APOA1 | COL1A1 | MMP12 | FPR1 | ANXA4 | HBA1/2 | MX1 |
| GPX3 | KITLG | SLPI | G0S2 | SLPI | ITGB8 | TOP2A | EPHB4 |
| SERPINA3 | ATP5F1B | SPINK1 | HSPA1A/B | CLEC4E | KRT7 | VHL | LYZ |
| LCN2 | RGS5 | THBS2 | HBA1/2 | KRT16 | CDH1 | ITGA6 | S100A9 |
| AZGP1 | CPA3 | CEACAM6 | PTGES | KRT17 | SLPI | UBE2C | MRC1 |
| TPT1 | BTG1 | KRT7 | HEY1 | | CCND1 | HSP90AB1 | HLA-DRA |
| SLC40A1 | CNTFR | KRT19 | LGR5 | | LGALS3 | FGF13 | G6PD |
| FABP4 | ST6GAL1 | SPP1 | VHL | | SERPINB5 | SST | HLA-DPB1 |
| CD36 | TPSAB1/B2 | TIMP1 | STMN1 | | EPCAM | MXRA8 | CAV1 |
| ZBTB16 | ACTA2 | COL3A1 | IGHG1 | | PTK6 | FASN | CD52 |
| DPP4 | KIT | MGP | MXRA8 | | DDR1 | SRC | HLA-DQA1 |
| VEGFD | COL6A1 | KRT23 | SST | | SPP1 | BMP4 | C1QA |
| FGG | TPT1 | CXCL5 | IGKC | | S100P | TUBB4B | DUSP5 |
| CTSD | VTN | | TNNC1 | | CD24 | TPM2 | ANXA2 |
| NCAM1 | HSP90B1 | | PTGDR2 | | KRT8 | NRXN3 | AIF1 |
| THSD4 | GAS6 | | ENO1 | | DCN | RNF43 | THBS1 |
| AR | LIFR | | SLC2A1 | | TIMP1 | EZH2 | OAS1 |
| SERPINA1 | CTSG | | PRF1 | | LUM | HSPA1A/B | HLA-DRB |
| APOC1 | EGFR | | COL14A1 | | TM4SF1 | PTK2 | CXCL8 |
| KRT20 | COL6A2 | | LAG3 | | HSD17B2 | CHEK2 | CFD |
| IL1R1 | ITGA9 | | LY6D | | CRYAB | TNFRSF19 | IL10RA |
| PIGR | SEC61G | | CD8A | | | SLC2A1 | CLEC5A |
| APOE | ITGA5 | | MEG3 | | | ANGPT1 | OLR1 |
| LAMP2 | CD34 | | MMP9 | | | TOX | CIITA |
| RORA | APOE | | BMP4 | | | VWA1 | MYC |
| KDR | NOTCH3 | | VWA1 | | | MAPK14 | LMNA |
| HMGCS1 | RACK1 | | LINC02446 | | | MEG3 | C1QC |
| | FKBP11 | | DMBT1 | | | BMP7 | LCN2 |
| | VIM | | XKR4 | | | HEY1 | TNFRSF12A |
| | | | ANGPT1 | | | GSTP1 | ISG15 |
| | | | FZD1 | | | MARCKSL1 | OAS3 |
| | | | WNT7A | | | ERBB3 | ALOX5AP |
| | | | CHEK2 | | | PTGES | MSR1 |
| | | | IGHA1 | | | PDGFA | CD4 |
| | | | CST7 | | | DNMT3A | SPP1 |
| | | | IGHM | | | FCER1G | TGFBI |
| | | | ACKR4 | | | HMGB2 | KRT80 |
| | | | RAG1 | | | HDAC4 | IFI44L |
| | | | RNF43 | | | HSPB1 | CSF2RA |
| | | | SNAI2 | | | HDAC11 | GPR183 |
| | | | TNXA/B | | | SQLE | ARHGDIB |
| | | | GZMK | | | ALCAM | ITGAM |

| village_1 | village_2 | village_3 | village_4 | village_5 | village_6 | village_7 | village_8 |
|---|---|---|---|---|---|---|---|
| | | | UBE2C | | | H4C3 | BST2 |
| | | | LINC01857 | | | PTGDR2 | PTGS1 |
| | | | CCR10 | | | RAMP1 | NOTCH2 |
| | | | GPER1 | | | TNNC1 | ITGA3 |
| | | | RAMP1 | | | PLCG1 | TPM1 |
| | | | GZMA | | | MMP12 | AXL |
| | | | PDCD1 | | | CENPF | CD44 |
| | | | SRC | | | GDF15 | CD5L |
| | | | CD27 | | | PCNA | CD68 |
| | | | CELSR2 | | | COL14A1 | IL17RA |
| | | | HDAC11 | | | HDAC1 | DDX58 |
| | | | GPBAR1 | | | HSP90AA1 | CENPF |
| | | | CCL26 | | | TYMS | HLA-DPA1 |
| | | | TNFRSF19 | | | CELSR1 | IFIT3 |
| | | | DNMT3A | | | GPER1 | H2AZ1 |
| | | | BEST1 | | | BBLN | MMP14 |
| | | | CALB1 | | | MIF | MB |
| | | | PTK2 | | | DDC | RPS4Y1 |
| | | | NELL2 | | | NR1H3 | NFKB1 |
| | | | FGF13 | | | PRF1 | CCL4/L1/L2 |
| | | | IFNL2/3 | | | DUSP6 | MS4A6A |
| | | | TCAP | | | SERPINH1 | PLAC8 |
| | | | IL7 | | | MMP2 | SMAD3 |
| | | | TPM2 | | | MYC | LGALS9 |
| | | | GZMB | | | GZMA | KRT15 |
| | | | FZD6 | | | MKI67 | ADGRE5 |
| | | | ALCAM | | | TGFB2 | AGR2 |
| | | | SNAI1 | | | RPS4Y1 | CRYAB |
| | | | RPS4Y1 | | | LDHA | IL2RB |
| | | | ADM2 | | | XKR4 | AZGP1 |
| | | | CD83 | | | CD8A | CD22 |
| | | | TGFB2 | | | SMARCB1 | LDHA |
| | | | HSP90AB1 | | | ETV5 | SQSTM1 |
| | | | PDGFA | | | CHEK1 | IFNGR1 |
| | | | CCR7 | | | BGN | CD47 |
| | | | HDAC4 | | | TNXA/B | IL2RG |
| | | | SPOCK2 | | | CELSR2 | SLCO2B1 |
| | | | FASLG | | | GZMK | CD93 |
| | | | TOP2A | | | SNAI2 | FCGR3A/B |
| | | | CSF3 | | | UBA52 | CLEC12A |
| | | | WNT10B | | | BRCA1 | TUBB4B |
| | | | CXCR5 | | | | SH3BGRL3 |
| | | | REG1A | | | | TAP1 |
| | | | IGFBP6 | | | | S100P |
| | | | ITGA6 | | | | SOX9 |
| | | | APOD | | | | ADGRE2 |
| | | | EOMES | | | | HEXB |
| | | | NR1H3 | | | | |
| | | | NLRC4 | | | | |

| village_1 | village_2 | village_3 | village_4 | village_5 | village_6 | village_7 | village_8 |
|---|---|---|---|---|---|---|---|
| | | | CYP1B1 | | | | |
| | | | IL17A | | | | |
| | | | CELSR1 | | | | |
| | | | FGF12 | | | | |
| | | | JCHAIN | | | | |
| | | | EZH2 | | | | |
| | | | PRSS2 | | | | |

Table S5: Top gene pairs in each tumor cell village

| Village | CellType | Gene_pair | Village | CellType | Gene_pair |
|---|---|---|---|---|---|
| village 1 | Tumor - TAM | LYZ - SERPINA1 | village 2 | Tumor - TAM | APOA1 - APOE |
| village 1 | Tumor - TAM | LYZ - FN1 | village 2 | Tumor - TEC | APOA1 - APOE |
| village 1 | Tumor - TAM | SERPINA1 - LYZ | village 2 | Tumor - CAF | APOA1 - APOE |
| village 1 | Tumor - TEC | LYZ - SERPINA1 | village 2 | Tumor - TAM | APOA1 - PPIA |
| village 1 | Tumor - CAF | LYZ - SERPINA1 | village 2 | Tumor - TAM | APOA1 - ATP5F1B |
| village 1 | Tumor - TAM | SERPINA1 - FN1 | village 2 | Tumor - TAM | GC - APOE |
| village 1 | Tumor - TAM | LYZ - SOD2 | village 2 | Tumor - TAM | IL32 - B2M |
| village 1 | Tumor - TAM | SERPINA1 - SOD2 | village 2 | Tumor - TAM | APOA1 - SERPINA1 |
| village 1 | Tumor - TAM | GC - SERPINA1 | village 2 | Tumor - TEC | APOA1 - ATP5F1B |
| village 1 | Tumor - TAM | FN1 - SERPINA1 | village 2 | Tumor - CAF | APOA1 - PPIA |
| village 1 | Tumor - TAM | FN1 - LYZ | village 2 | Tumor - TAM | APOA1 - ATP5F1E |
| village 1 | Tumor - TEC | GC - SERPINA1 | village 2 | Tumor - CAF | GC - APOE |
| village 1 | Tumor - TAM | GC - FN1 | village 2 | Tumor - TEC | APOA1 - SERPINA1 |
| village 1 | Tumor - TAM | SERPINA1 - GLUL | village 2 | Tumor - CAF | APOA1 - ATP5F1B |
| village 1 | Tumor - TAM | LYZ - GLUL | village 2 | Tumor - TEC | GC - APOE |
| village 1 | Tumor - TEC | FN1 - SERPINA1 | village 2 | Tumor - TEC | IL32 - B2M |
| village 1 | Tumor - CAF | GC - SERPINA1 | village 2 | Tumor - CAF | IL32 - B2M |
| village 1 | Tumor - CAF | FN1 - SERPINA1 | village 2 | Tumor - CAF | APOA1 - SERPINA1 |
| village 1 | Tumor - TAM | LYZ - IFITM3 | village 2 | Tumor - TEC | APOA1 - PPIA |
| village 1 | Tumor - TEC | FGG - SERPINA1 | village 2 | Tumor - CAF | APOE - PPIA |
| village 1 | Tumor - TAM | GC - LYZ | village 2 | Tumor - TAM | APOA1 - FN1 |
| village 1 | Tumor - CAF | LYZ - GLUL | village 2 | Tumor - TAM | GC - SERPINA1 |
| village 1 | Tumor - TAM | LYZ - CTSD | village 2 | Tumor - TAM | GC - ATP5F1B |
| village 1 | Tumor - TAM | CTSD - SERPINA1 | village 2 | Tumor - TAM | PPIA - APOE |
| village 1 | Tumor - CAF | SERPINA1 - GLUL | village 2 | Tumor - TEC | APOA1 - ATP5F1E |
| village 1 | Tumor - TAM | SERPINA1 - CTSD | village 2 | Tumor - TEC | B2M - IL32 |
| village 1 | Tumor - TAM | CTSD - LYZ | village 2 | Tumor - TAM | ATP5F1B - APOE |
| village 1 | Tumor - TAM | FGG - SERPINA1 | village 2 | Tumor - TAM | APOA1 - CD63 |
| village 1 | Tumor - TAM | SERPINA1 - IFITM3 | village 2 | Tumor - TAM | GC - ATP5F1E |
| village 1 | Tumor - TAM | B2M - SERPINA1 | village 2 | Tumor - CAF | APOA1 - ATP5F1E |
| village 1 | Tumor - TAM | CLU - SERPINA1 | village 2 | Tumor - TAM | ATP5F1B - PPIA |
| village 1 | Tumor - TAM | LYZ - LAMP2 | village 2 | Tumor - TAM | PPIA - ATP5F1B |
| village 1 | Tumor - CAF | SERPINA1 - TIMP1 | village 2 | Tumor - TAM | APOA1 - CTNNB1 |
| village 1 | Tumor - TAM | B2M - LYZ | village 2 | Tumor - TAM | GC - PPIA |
| village 1 | Tumor - CAF | SERPINA1 - ITM2B | village 2 | Tumor - CAF | GC - SERPINA1 |
| village 1 | Tumor - TAM | TTR - GLUL | village 2 | Tumor - TAM | CTNNB1 - APOE |
| village 1 | Tumor - TAM | CTSD - FN1 | village 2 | Tumor - CAF | PPIA - APOE |
| village 1 | Tumor - TAM | SERPINA1 - LAMP2 | village 2 | Tumor - TEC | PPIA - APOE |
| village 1 | Tumor - TAM | B2M - FN1 | village 2 | Tumor - TEC | GC - SERPINA1 |
| village 1 | Tumor - TEC | CTSD - SERPINA1 | village 2 | Tumor - TAM | APOA1 - HMGN2 |
| village 1 | Tumor - TEC | SERPINA1 - GLUL | village 2 | Tumor - CAF | APOE - ATP5F1B |
| village 1 | Tumor - CAF | LYZ - ITM2B | village 2 | Tumor - TAM | VTN - APOE |
| village 1 | Tumor - CAF | CTSD - SERPINA1 | village 2 | Tumor - CAF | APOA1 - CTNNB1 |
| village 1 | Tumor - TAM | SERPINA1 - IL32 | village 2 | Tumor - CAF | GC - PPIA |
| village 1 | Tumor - CAF | APOE - SERPINA1 | village 2 | Tumor - TAM | CALM2 - PPIA |
| village 1 | Tumor - TAM | NEAT1 - TPT1 | village 2 | Tumor - TAM | APOA1 - HSP90B1 |
| village 1 | Tumor - TEC | LYZ - FN1 | village 2 | Tumor - CAF | ATP5F1B - PPIA |
| village 1 | Tumor - TEC | APOE - GLUL | village 2 | Tumor - TAM | B2M - CD74 |
| village 1 | Tumor - TAM | B2M - SOD2 | village 2 | Tumor - TEC | PPIA - ATP5F1B |
| village 1 | Tumor - CAF | SERPINA1 - IFITM3 | village 2 | Tumor - TAM | CALM2 - APOE |

| Village | CellType | Gene_pair | Village | CellType | Gene_pair |
|---|---|---|---|---|---|
| village 3 | Tumor - CAF | S100A6 - MGP | village 4 | Tumor - CAF | APOE - APOC1 |
| village 3 | Tumor - TAM | APOE - SERPINA1 | village 4 | Tumor - TAM | APOA1 - SERPINA1 |
| village 3 | Tumor - TAM | SPP1 - SERPINA1 | village 4 | Tumor - TAM | APOE - APOC1 |
| village 3 | Tumor - CAF | S100A6 - TIMP1 | village 4 | Tumor - T-cell | APOC1 - MALAT1 |
| village 3 | Tumor - TAM | CRP - SPP1 | village 4 | Tumor - T-cell | NEAT1 - MALAT1 |
| village 3 | Tumor - CAF | SERPINA1 - COL6A1 | village 4 | Tumor - T-cell | MALAT1 - NEAT1 |
| village 3 | Tumor - CAF | SPINK1 - BGN | village 4 | Tumor - CAF | NEAT1 - MALAT1 |
| village 3 | Tumor - CAF | SERPINA1 - B2M | village 4 | Tumor - CAF | MALAT1 - NEAT1 |
| village 3 | Tumor - CAF | RARRES2 - COL6A1 | village 4 | Tumor - TAM | APOC1 - B2M |
| village 3 | Tumor - CAF | TPT1 - BGN | village 4 | Tumor - T-cell | HSP90AA1 - HSPB1 |
| village 3 | Tumor - CAF | LYZ - BGN | village 4 | Tumor - TAM | MALAT1 - NEAT1 |
| village 3 | Tumor - TAM | RARRES2 - SERPINA1 | village 4 | Tumor - TAM | APOC1 - CD74 |
| village 3 | Tumor - TAM | SOD2 - SPP1 | village 4 | Tumor - T-cell | APOC1 - B2M |
| village 3 | Tumor - TAM | SERPINA1 - GLUL | village 4 | Tumor - T-cell | APOE - MALAT1 |
| village 3 | Tumor - CAF | ITM2B - BGN | village 4 | Tumor - CAF | HSPB1 - HSP90AA1 |
| village 3 | Tumor - TAM | S100A6 - IFI6 | village 4 | Tumor - TAM | APOC1 - MIF |
| village 3 | Tumor - TAM | LYZ - IGFBP7 | village 4 | Tumor - TAM | B2M - CD74 |
| village 3 | Tumor - CAF | SERPINA1 - MT2A | village 4 | Tumor - T-cell | HSPB1 - MALAT1 |
| village 3 | Tumor - TAM | SERPINA1 - IL32 | village 4 | Tumor - TAM | APOC1 - TPT1 |
| village 3 | Tumor - CAF | VTN - COL6A1 | village 4 | Tumor - CAF | APOC1 - B2M |
| village 3 | Tumor - CAF | SPP1 - SERPINA1 | village 4 | Tumor - CAF | HSP90AA1 - HSPB1 |
| village 3 | Tumor - CAF | CRYAB - IFI6 | village 4 | Tumor - TAM | UBA52 - PFN1 |
| village 3 | Tumor - CAF | B2M - IFI6 | village 4 | Tumor - TAM | MIF - CD74 |
| village 3 | Tumor - CAF | RARRES2 - SERPINA1 | village 4 | Tumor - TAM | APOC1 - APOE |
| village 3 | Tumor - CAF | RARRES2 - B2M | village 4 | Tumor - TAM | HSPB1 - HSP90AA1 |
| village 3 | Tumor - CAF | MIF - TIMP1 | village 4 | Tumor - CAF | APOC1 - APOE |
| village 3 | Tumor - CAF | LYZ - VIM | village 4 | Tumor - TAM | MIF - UBA52 |
| village 3 | Tumor - CAF | PFN1 - TIMP1 | village 4 | Tumor - CAF | HSPB1 - MALAT1 |
| village 3 | Tumor - CAF | VTN - DCN | village 4 | Tumor - CAF | APOC1 - MIF |
| village 3 | Tumor - CAF | TPT1 - DCN | village 4 | Tumor - TAM | MIF - B2M |
| village 3 | Tumor - CAF | SPINK1 - COL5A1 | village 4 | Tumor - CAF | APOC1 - MALAT1 |
| village 3 | Tumor - CAF | S100A6 - IFI6 | village 4 | Tumor - TAM | PFN1 - CD74 |
| village 3 | Tumor - CAF | CRYAB - B2M | village 4 | Tumor - CAF | APOC1 - TPT1 |
| village 3 | Tumor - CAF | S100A6 - DCN | village 4 | Tumor - TAM | APOC1 - UBA52 |
| village 3 | Tumor - CAF | S100A10 - BGN | village 4 | Tumor - TAM | HSP90AA1 - HSPB1 |
| village 3 | Tumor - CAF | S100A6 - MT2A | village 4 | Tumor - T-cell | MALAT1 - HSPB1 |
| village 3 | Tumor - CAF | KRT19 - MGP | village 4 | Tumor - TAM | PFN1 - B2M |
| village 3 | Tumor - CAF | TPT1 - COL1A1 | village 4 | Tumor - CAF | UBA52 - MIF |
| village 3 | Tumor - TAM | B2M - IFI6 | village 4 | Tumor - T-cell | HSPB1 - HSP90AA1 |
| village 3 | Tumor - CAF | TPT1 - TIMP1 | village 4 | Tumor - TAM | MIF - PFN1 |
| village 3 | Tumor - TAM | CRYAB - IFI6 | village 4 | Tumor - TAM | NEAT1 - MALAT1 |
| village 3 | Tumor - CAF | S100A10 - TIMP1 | village 4 | Tumor - CAF | APOE - MIF |
| village 3 | Tumor - TAM | MMP7 - SPP1 | village 4 | Tumor - CAF | APOE - B2M |
| village 3 | Tumor - CAF | S100A6 - COL1A1 | village 4 | Tumor - CAF | MIF - PFN1 |
| village 3 | Tumor - CAF | VTN - TIMP1 | village 4 | Tumor - TAM | APOE - MIF |
| village 3 | Tumor - CAF | S100A6 - TPT1 | village 4 | Tumor - TAM | MIF - APOC1 |
| village 3 | Tumor - TAM | ITM2B - IGFBP7 | village 4 | Tumor - T-cell | APOC1 - TPT1 |
| village 3 | Tumor - CAF | LYZ - COL5A1 | village 4 | Tumor - TAM | UBA52 - MIF |
| village 3 | Tumor - CAF | CRYAB - MT2A | village 4 | Tumor - CAF | MALAT1 - APOC1 |
| village 3 | Tumor - CAF | TPT1 - COL5A1 | village 4 | Tumor - TAM | TPT1 - UBA52 |

| Village | CellType | Gene_pair | Village | CellType | Gene_pair |
|---|---|---|---|---|---|
| village 5 | Tumor - TAM | SPP1 - SERPINA1 | village 6 | Tumor - CAF | SPINK1 - BGN |
| village 5 | Tumor - CAF | SPP1 - COL3A1 | village 6 | Tumor - TAM | SPINK1 - IGFBP7 |
| village 5 | Tumor - TAM | SPP1 - LYZ | village 6 | Tumor - CAF | ITM2B - BGN |
| village 5 | Tumor - CAF | SPP1 - FN1 | village 6 | Tumor - CAF | SPINK1 - COL5A1 |
| village 5 | Tumor - CAF | SERPINA1 - COL1A1 | village 6 | Tumor - CAF | LYZ - BGN |
| village 5 | Tumor - CAF | SPP1 - COL1A1 | village 6 | Tumor - CAF | SPINK1 - COL1A2 |
| village 5 | Tumor - CAF | VTN - IGFBP7 | village 6 | Tumor - TAM | SPP1 - SERPINA1 |
| village 5 | Tumor - CAF | SERPINA1 - TIMP1 | village 6 | Tumor - CAF | SPINK1 - VIM |
| village 5 | Tumor - CAF | SERPINA1 - COL3A1 | village 6 | Tumor - TAM | SPP1 - B2M |
| village 5 | Tumor - CAF | SPP1 - IFI6 | village 6 | Tumor - TAM | ITM2B - IGFBP7 |
| village 5 | Tumor - CAF | SPP1 - TIMP1 | village 6 | Tumor - TAM | VTN - SERPINA1 |
| village 5 | Tumor - TAM | SPP1 - HLA - DRA | village 6 | Tumor - CAF | SPINK1 - COL1A1 |
| village 5 | Tumor - CAF | IFI6 - COL1A1 | village 6 | Tumor - CAF | HSPB1 - BGN |
| village 5 | Tumor - CAF | CRYAB - IFI6 | village 6 | Tumor - TAM | SPINK1 - VIM |
| village 5 | Tumor - CAF | SERPINA1 - FN1 | village 6 | Tumor - CAF | TPT1 - BGN |
| village 5 | Tumor - TAM | SERPINA1 - HLA - DRA | village 6 | Tumor - TAM | SPP1 - IFI6 |
| village 5 | Tumor - TAM | SPP1 - GLUL | village 6 | Tumor - CAF | SPINK1 - COL4A1 |
| village 5 | Tumor - TAM | SPP1 - MT2A | village 6 | Tumor - TAM | SPP1 - C1QA |
| village 5 | Tumor - CAF | VTN - RGS5 | village 6 | Tumor - CAF | ITM2B - COL5A1 |
| village 5 | Tumor - CAF | IFI6 - IFITM1 | village 6 | Tumor - CAF | VTN - SERPINA1 |
| village 5 | Tumor - CAF | B2M - MGP | village 6 | Tumor - TAM | VTN - PPIA |
| village 5 | Tumor - CAF | PFN1 - COL1A1 | village 6 | Tumor - TAM | CLU - HLA - DRA |
| village 5 | Tumor - CAF | CCND1 - IGFBP7 | village 6 | Tumor - CAF | SPINK1 - IGFBP7 |
| village 5 | Tumor - TAM | SPP1 - CD74 | village 6 | Tumor - CAF | SPINK1 - MGP |
| village 5 | Tumor - CAF | SPP1 - COL6A3 | village 6 | Tumor - CAF | VEGFA - BGN |
| village 5 | Tumor - TAM | SPP1 - C1QC | village 6 | Tumor - CAF | LYZ - IGFBP7 |
| village 5 | Tumor - CAF | SPP1 - LGALS1 | village 6 | Tumor - TAM | CLU - C1QC |
| village 5 | Tumor - TAM | SERPINA1 - HLA - DPA1 | village 6 | Tumor - TAM | TPT1 - IGFBP7 |
| village 5 | Tumor - TAM | SPP1 - HLA - DPA1 | village 6 | Tumor - CAF | CRYAB - RGS5 |
| village 5 | Tumor - CAF | SPP1 - IFITM1 | village 6 | Tumor - TAM | SERPINA1 - PPIA |
| village 5 | Tumor - TAM | SERPINA1 - LYZ | village 6 | Tumor - CAF | LYZ - LUM |
| village 5 | Tumor - CAF | SLPI - RGS5 | village 6 | Tumor - CAF | ITM2B - COL1A2 |
| village 5 | Tumor - TAM | CRYAB - IFI6 | village 6 | Tumor - TAM | SPP1 - C1QB |
| village 5 | Tumor - CAF | SLPI - APOE | village 6 | Tumor - CAF | SPP1 - SERPINA1 |
| village 5 | Tumor - CAF | SPP1 - IFITM3 | village 6 | Tumor - TAM | SPP1 - GLUL |
| village 5 | Tumor - CAF | VTN - APOE | village 6 | Tumor - TAM | SERPINA1 - HLA - DRA |
| village 5 | Tumor - CAF | IFI6 - TIMP1 | village 6 | Tumor - CAF | ITM2B - COL1A1 |
| village 5 | Tumor - CAF | MIF - COL1A1 | village 6 | Tumor - CAF | SPINK1 - TPT1 |
| village 5 | Tumor - TAM | SERPINA1 - C1QC | village 6 | Tumor - TAM | SPP1 - HLA - DRA |
| village 5 | Tumor - CAF | REG1A - TAGLN | village 6 | Tumor - CAF | ITM2B - LUM |
| village 5 | Tumor - CAF | KRT19 - IGFBP7 | village 6 | Tumor - TAM | VTN - HLA - DRA |
| village 5 | Tumor - CAF | B2M - IGFBP7 | village 6 | Tumor - CAF | SPP1 - COL6A1 |
| village 5 | Tumor - CAF | SPP1 - COL4A1 | village 6 | Tumor - TAM | HSPB1 - IGFBP7 |
| village 5 | Tumor - CAF | REG1A - IGFBP7 | village 6 | Tumor - CAF | ITM2B - IGFBP7 |
| village 5 | Tumor - CAF | PFN1 - TIMP1 | village 6 | Tumor - CAF | SPINK1 - LUM |
| village 5 | Tumor - TAM | SERPINA1 - SPP1 | village 6 | Tumor - TAM | SPP1 - C1QC |
| village 5 | Tumor - TAM | SPP1 - IFI6 | village 6 | Tumor - CAF | SERPINA1 - COL6A1 |
| village 5 | Tumor - CAF | ANXA4 - IGFBP7 | village 6 | Tumor - CAF | LYZ - COL1A1 |
| village 5 | Tumor - CAF | SPP1 - B2M | village 6 | Tumor - TAM | STAT1 - HLA - DRA |
| village 5 | Tumor - CAF | SPP1 - DCN | village 6 | Tumor - CAF | NEAT1 - MALAT1 |

| Village | CellType | Gene_pair | Village | CellType | Gene_pair |
|---|---|---|---|---|---|
| village 7 | Tumor - TAM | HSP90B1 - B2M | village 8 | Tumor - CAF | SOD2 - COL1A1 |
| village 7 | Tumor - TAM | APOC1 - SERPINA1 | village 8 | Tumor - CAF | S100A6 - COL1A1 |
| village 7 | Tumor - TAM | APOA1 - SERPINA1 | village 8 | Tumor - TAM | S100A6 - APOE |
| village 7 | Tumor - TAM | APOE - SERPINA1 | village 8 | Tumor - T-cell | MET - ANXA2 |
| village 7 | Tumor - TAM | UBA52 - MIF | village 8 | Tumor - T-cell | APOC1 - SQSTM1 |
| village 7 | Tumor - TAM | SAT1 - B2M | village 8 | Tumor - CAF | ANXA2 - B2M |
| village 7 | Tumor - TAM | GSTP1 - C1QC | village 8 | Tumor - T-cell | APOC1 - ANXA2 |
| village 7 | Tumor - TAM | SAT1 - C1QC | village 8 | Tumor - CAF | MET - VIM |
| village 7 | Tumor - TAM | BGN - C1QC | village 8 | Tumor - TAM | APOC1 - LYZ |
| village 7 | Tumor - TAM | SAT1 - C1QB | village 8 | Tumor - CAF | ANXA2 - VIM |
| village 7 | Tumor - TAM | GLUL - APOE | village 8 | Tumor - CAF | MET - B2M |
| village 7 | Tumor - TAM | APOE - GLUL | village 8 | Tumor - TAM | MET - ANXA2 |
| village 7 | Tumor - TAM | HSP90B1 - C1QC | village 8 | Tumor - TAM | APOC1 - HLA - DRA |
| village 7 | Tumor - TAM | BGN - PFN1 | village 8 | Tumor - CAF | APOC1 - VIM |
| village 7 | Tumor - TAM | HSP90AB1 - PFN1 | village 8 | Tumor - CAF | APOC1 - COL6A2 |
| village 7 | Tumor - TAM | MALAT1 - NEAT1 | village 8 | Tumor - CAF | MET - ITGB1 |
| village 7 | Tumor - TAM | GSTP1 - B2M | village 8 | Tumor - CAF | MET - COL1A2 |
| village 7 | Tumor - TAM | GSTP1 - C1QB | village 8 | Tumor - TAM | APOC1 - SQSTM1 |
| village 7 | Tumor - TAM | APOE - PSAP | village 8 | Tumor - CAF | ANXA2 - COL1A2 |
| village 7 | Tumor - TAM | SERPINA1 - UBA52 | village 8 | Tumor - T-cell | MET - SQSTM1 |
| village 7 | Tumor - TAM | BGN - B2M | village 8 | Tumor - CAF | APOC1 - B2M |
| village 7 | Tumor - TAM | BGN - C1QB | village 8 | Tumor - TAM | APOC1 - MYL12A |
| village 7 | Tumor - TAM | GLUL - B2M | village 8 | Tumor - TAM | APOC1 - ANXA2 |
| village 7 | Tumor - TAM | GSTP1 - CD74 | village 8 | Tumor - TAM | MET - LYZ |
| village 7 | Tumor - TAM | ENO1 - SPP1 | village 8 | Tumor - CAF | APOC1 - COL1A2 |
| village 7 | Tumor - TAM | TUBB4B - C1QB | village 8 | Tumor - CAF | APOC1 - ITGB1 |
| village 7 | Tumor - TAM | GLUL - CD74 | village 8 | Tumor - CAF | LYZ - COL6A2 |
| village 7 | Tumor - TAM | HSP90B1 - CD74 | village 8 | Tumor - T-cell | FN1 - ANXA2 |
| village 7 | Tumor - TAM | SAT1 - CD74 | village 8 | Tumor - TAM | MET - HLA - DRA |
| village 7 | Tumor - TAM | BGN - CD74 | village 8 | Tumor - T-cell | APOC1 - MYL12A |
| village 7 | Tumor - TAM | GLUL - PSAP | village 8 | Tumor - TAM | EPHB4 - LYZ |
| village 7 | Tumor - TAM | APOC1 - LYZ | village 8 | Tumor - CAF | SOD2 - MGP |
| village 7 | Tumor - TAM | ATP5F1E - PFN1 | village 8 | Tumor - CAF | SOD2 - S100A6 |
| village 7 | Tumor - TAM | B2M - CD74 | village 8 | Tumor - CAF | SQSTM1 - VIM |
| village 7 | Tumor - TAM | LDHA - ENO1 | village 8 | Tumor - TAM | APOC1 - FCGBP |
| village 7 | Tumor - TAM | HSP90B1 - PFN1 | village 8 | Tumor - T-cell | FGG - ANXA2 |
| village 7 | Tumor - TAM | SQSTM1 - GLUL | village 8 | Tumor - CAF | S100A6 - MGP |
| village 7 | Tumor - TAM | GLUL - C1QB | village 8 | Tumor - T-cell | CAV1 - ANXA2 |
| village 7 | Tumor - TAM | UBA52 - HMGN2 | village 8 | Tumor - TAM | FGG - LYZ |
| village 7 | Tumor - TAM | GLUL - HLA - DRA | village 8 | Tumor - TAM | NDRG1 - SPP1 |
| village 7 | Tumor - TAM | IFITM3 - B2M | village 8 | Tumor - TAM | APOC1 - C1QA |
| village 7 | Tumor - TAM | TUBB4B - C1QC | village 8 | Tumor - CAF | CAV1 - ANXA2 |
| village 7 | Tumor - TAM | TPT1 - UBA52 | village 8 | Tumor - T-cell | SQSTM1 - ANXA2 |
| village 7 | Tumor - TAM | HSP90B1 - C1QB | village 8 | Tumor - CAF | ANXA2 - ITGB1 |
| village 7 | Tumor - TAM | APOE - LYZ | village 8 | Tumor - TAM | FN1 - LYZ |
| village 7 | Tumor - TAM | GLUL - HLA - DQA1 | village 8 | Tumor - CAF | MET - ANXA2 |
| village 7 | Tumor - TAM | B2M - HLA - DRA | village 8 | Tumor - T-cell | AZGP1 - ANXA2 |
| village 7 | Tumor - TAM | GSTP1 - PFN1 | village 8 | Tumor - CAF | SQSTM1 - B2M |
| village 7 | Tumor - TAM | UBA52 - HSP90AA1 | village 8 | Tumor - CAF | APOC1 - MYL12A |
| village 7 | Tumor - TAM | TPT1 - ATP5F1E | village 8 | Tumor - TAM | CAV1 - ANXA2 |
